# Information Theoretic Feature Selection Methods for Single Cell RNA-Sequencing

**DOI:** 10.1101/646919

**Authors:** Umang Varma, Justin Colacino, Anna Gilbert

## Abstract

Single cell RNA-sequencing (scRNA-seq) technologies have generated an expansive amount of new biological information, revealing new cellular populations and hierarchical relationships. A number of technologies complementary to scRNA-seq rely on the selection of a smaller number of marker genes (or features) to accurately differentiate cell types within a complex mixture of cells. In this paper, we benchmark differential expression methods against information-theoretic feature selection methods to evaluate the ability of these algorithms to identify small and efficient sets of genes that are informative about cell types. Unlike differential methods, that are strictly binary and univariate, information-theoretic methods can be used as any combination of binary or multiclass and univariate or multivariate. We show for some datasets, information theoretic methods can reveal genes that are both distinct from those selected by traditional algorithms and that are as informative, if not more, of the class labels. We also present detailed and principled theoretical analyses of these algorithms. All information theoretic methods in this paper are implemented in our PicturedRocks Python package that is compatible with the widely used scanpy package.

## 1 Background

Single cell RNA-sequencing (scRNA-seq) technologies have generated an expansive amount of new biological information, revealing new cellular populations and hierarchical relationships. These methods quantify expression of thousands of genes across tens or hundreds of thousands of individual cells derived from a single cell suspension [1, 2]. Using unbiased methods for cell clustering, researchers define cell types based on these gene expression patterns [3]. When information derived from these transcriptionally-defined cell clusters is integrated with spatial or protein level data, additional biological insights can be gleaned. A number of technologies complementary to scRNA-seq rely on the selection of a smaller number of marker genes to accurately differentiate cell types within a complex mixture of cells. For example, fluorescence activated cell sorting (FACS) can isolate specific cell populations from a single cell suspension for further in depth biological characterization based on the expression of a distinguishing set of a small number of cell surface markers. Mass cytometry provides data on expression of up to 40 proteins for cells in a single cell suspension, adapting traditional flow cytometry methods by labeling antibodies with heavy metal isotopes rather than fluorophores and using a time-of-flight mass spectrometry readout [4]. Imaging mass cytometry further allows for in situ assessment of approximately 40 markers in tissue sections, quantifying the spatial distribution of marker expression in the tissue [5]. Finally, spatial transcriptomic profiling approaches use fluorescence in situ hybridization in tissue sections for imaging based quantification of gene expression [6, 7]. Optimal usage of each of these techniques requires selection of a small set of marker genes (or features) that can accurately assign an individual cell to a known cell type based on expression.

Feature selection is a challenging combinatorial problem; it is already known that feature selection is NP-hard [8]. In the field of scRNA-seq, the most commonly used techniques for marker gene selection data are based on highly variable gene identification [9, 10], differential expression, and logistic regression [11], the former being unsupervised and the later supervised. These methods are univariate: they consider properties of each feature individually and do not consider any interactions between features. Additionally, these methods make strong assumptions about how gene expression is distributed. They typically measure the mean and standard deviation of features and use these in their determination of variability (or differential expression). Such techniques, especially ones that output probability values, are implicitly or explicitly making assumptions about the underlying distributions (e.g., Gaussian, or Student’s t-distribution). Even when the data suggest that these assumptions are reasonable, it is usually only the case for individual features in isolation. Supervised feature selection methods used in scRNAseq such as differential expression can natively handle binary class labels and need strategies such as one-vs-rest (OvR) or one-vs-one (OvO) to transform the problem back into a binary classification problem. This transformation causes two additional problems. First, by pooling clusters together (e.g., with OvR), we blur out valuable information by artificially suppressing genes that are informative about multiple class labels and boosting genes that are differentially expressed in one class when compared to all other classes. This puts an unnecessary importance on choosing class labels with the correct resolution. Secondly, methods to combining the lists of features identified in each pairwise comparison (either OvO or OvR) tend to be arbitrary and there is no canonical approach to reconciling the many (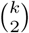 or *k* − 1 in OvO and OvR respectively, where *k* is the number of class labels) lists of features into one list.

In contrast to differential expression based methods for marker selection, information-theoretic methods address a number of these issues. Rather than considering properties of each marker individually, information-theoretic algorithms consider interactions between features to various degrees. Specifically, the “degree *K*” algorithm considers interactions between *K* features. For example, the degree 1 algorithm is a univariate algorithm, much like the differential expression methods, and the degree 2 algorithm considers interactions between pairs of features, etc. Additionally, in contrast to differential expression methods, information-theoretic methods can run as native multiclass methods as well as binary methods.

Our goal is to benchmark the efficacy of differential expression and information-theoretic methods for marker selection in scRNA-seq experiments, quantifying the strengths and weaknesses of a range of objective functions. We then focus on the degree 2 approximation, which corresponds to the conditional infomax feature extraction (CIFE) feature selection algorithm [12], and give theoretical guarantees for its performance. We then evaluate the performance of these algorithms on publicly available scRNA-seq datasets and describe optimizations for sparse data that improve the computational efficiency of these algorithms considerably.

## 2 Results

To assess the performance of mutual information methods in practice, we performed numerous experiments with publicly available single cell RNA-seq datasets. For each of the datasets listed below, we treat each cell as an observation and each gene as a feature.

- Paul dataset from [13] with 2730 mouse bone marrow cells and 3451 genes. Bone marrow cells form a continuous trajectory rather than distinct clusters because many of the cells are in the process of differentiating. The authors of the study assign class labels to cells based on expression of known marker genes.
- Green dataset [14] with 34633 cells from adult mouse testis and 37241 genes. The cells in dataset form a continuous trajectory, as should be expected for spermatogenesis.
- Zheng dataset from [10] with 68579 human blood cells and 32738 genes. This dataset consists of human blood cells that were clustered into distinct cell types. Zheng, et alt?hen used these clusters to isolate more cells based on these types to validate their computational results.
- Zeisel dataset [15] with 3005 mouse neuron cells and 19972 genes. Because neuron cells lie in discrete clusters in gene space, we expect clustering and classification tasks to be easier for this dataset. Further, Zeisel, et al. have carefully assigned labels to clusters and validated their cluster labels as corresponding to known biology.

Recall that our goal is to find a small set of genes that explain the given target labels (e.g., cluster labels or biologically verified cell type labels). To this end, we performed 5-fold cross validation by splitting the datasets into 5 folds, and then, for each fold, we selected markers on the four remaining folds, projected the data onto the selected features (i.e., only considered the columns of the matrix that correspond to the selected genes), trained a classifier on the four folds, and predicted class labels on the current fold. The feature selection methods that perform well are those that select discriminative features that, together, help predict the class of an unseen observation. In other words, feature selection methods with the lowest error rates choose genes that explain the class labels best.

We tested three mutual information based feature selection algorithms:

- the CIFE algorithm (which is equivalent to the degree 2 approximation presented under Degree K Supervised Feature Selection Algorithm on page 20),
- the MIM algorithm (which is equivalent to the degree 1 approximation), and
- the JMI algorithm (which dampens the redundancy penalty in CIFE by a factor of 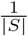, where *S* is the set of previously selected features).

For each dataset, we ran these algorithms in both a multi-class and binary fashion (see Multiclass vs Binary Feature Selection Methods). For all information-theoretic algorithms, we began by selecting the 5000 most relevant features (as found by the univariate MIM) before running bivariate algorithms (the run times in Figure 11 includes the MIM run time). For binary versions, this univariate filter was run for each class label separately.

We tested our algorithms with two different classifiers. Note that even for the datasets where class labels were originally found using an (unsupervised) classifier (rather than biologically), and we use supervised classifiers because we want to assess how informative the selected features are about the class labels.

- a Nearest Centroids Classifier that computes the centroid (unweighted mean of vectors) for each class label and to each unseen observation assigns the class label with the nearest centroid. We ran our experiments with this classifier to evaluate performance with a transparent classifier with minimal bells and whistles that can be easily implemented from scratch. Plots for classification errors with Nearest Centroids in the Figures section at the end of this paper.
- a Random Forests Classifier that uses an ensemble of decision trees. We use the Scikit Learn ([16]) implementation, which has numerous tweaks to improve accuracy and reduce overfitting. Although it is a more opaque classifier, the Random Forests Classifier does not give each feature equal weight and will use new features only when they are helpful. This aligns more closely with our motivation of finding an efficient set of features that are collective informative about the class labels rather than features that, individually, distinguish classes from one another.

We also tested against three differential expression algorithms (*t*-test with overestimated variation, Wilcoxon Rank Sum, and Logistic Regression) for single cell RNA sequencing (using the implementations in the widely used scanpy package^1^, see [17]). Since differential expression algorithms take a One-vs-Rest approach, we combined the lists into a single list by taking the first *k* genes (features) for each class label and removing duplicates.

Finally, since we are using a Random Forests classifier, we also evaluated Random Forests’ “feature importance” as a feature selection method. We should note that using Random Forests for feature selection and comparing against other methods using Random Forests is not a fair comparison, but we include it nonetheless for sake of comparison.

Figures 1, 2, 3, 4, 5, 6 are *t*-SNE plots (the projection of the dataset onto the top two principal components) for the Green dataset [14] generated using only a few genes. The colors of each point represents its class label (cell type).

**Figure 1:**
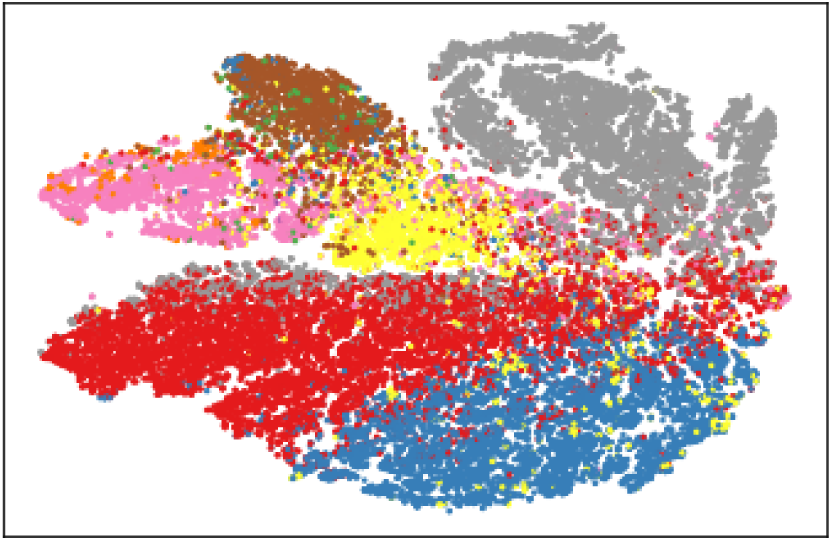
*t*-SNE Plot for Green Dataset using 10 features selected by CIFE (multiclass)

**Figure 2:**
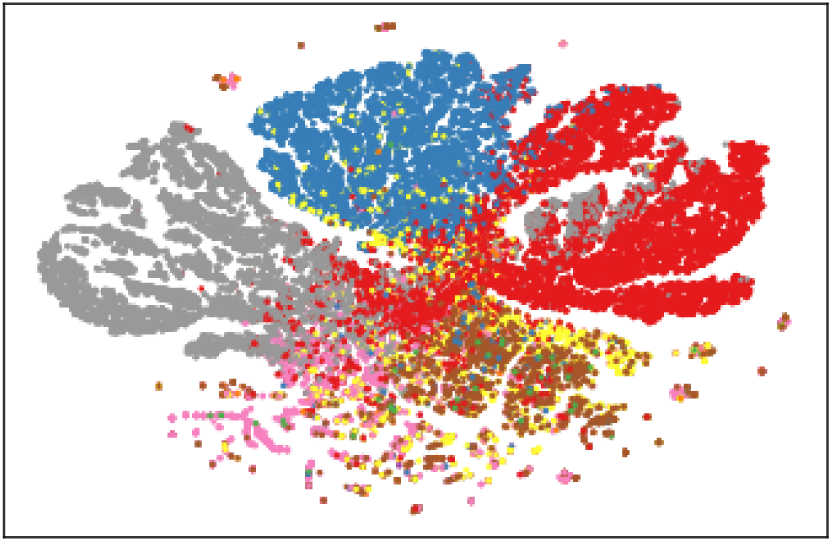
*t*-SNE Plot for Green Dataset using 10 features selected by JMI (multiclass)

**Figure 3:**
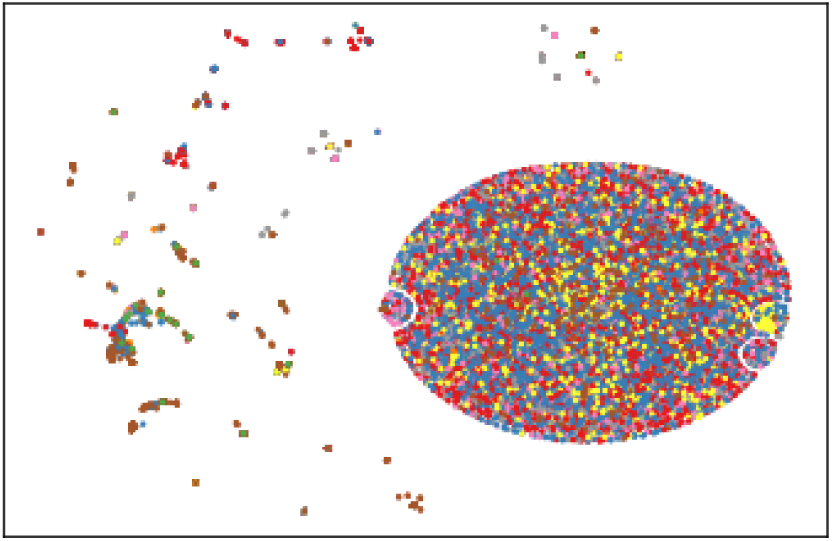
*t*-SNE Plot for Green Dataset using 11 features selected by Logistic Regression

**Figure 4:**
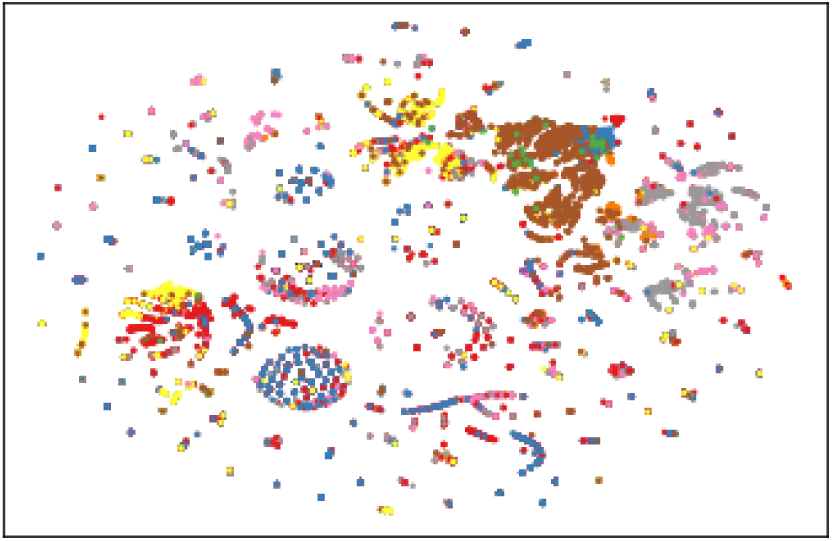
*t*-SNE Plot for Green Dataset using 11 features selected by *t*-test

**Figure 5:**
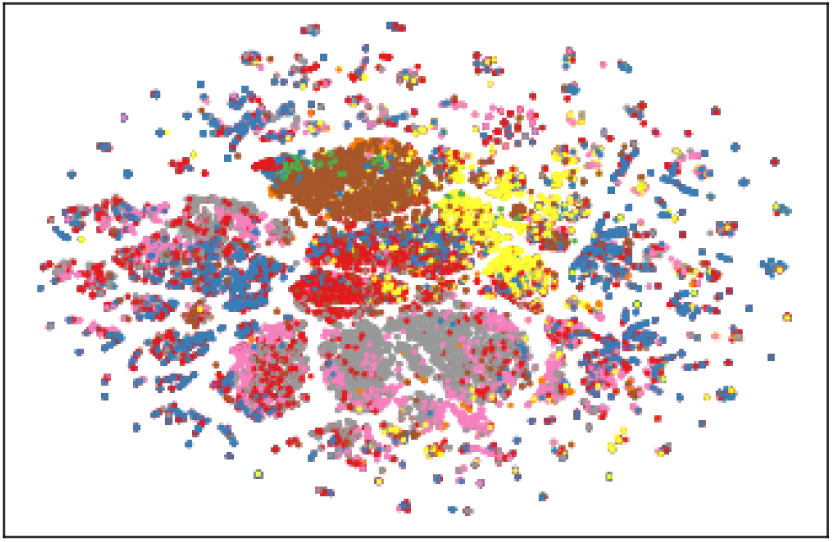
*t*-SNE Plot for Green Dataset using 11 features selected by Wilcoxon Rank Sum

**Figure 6:**
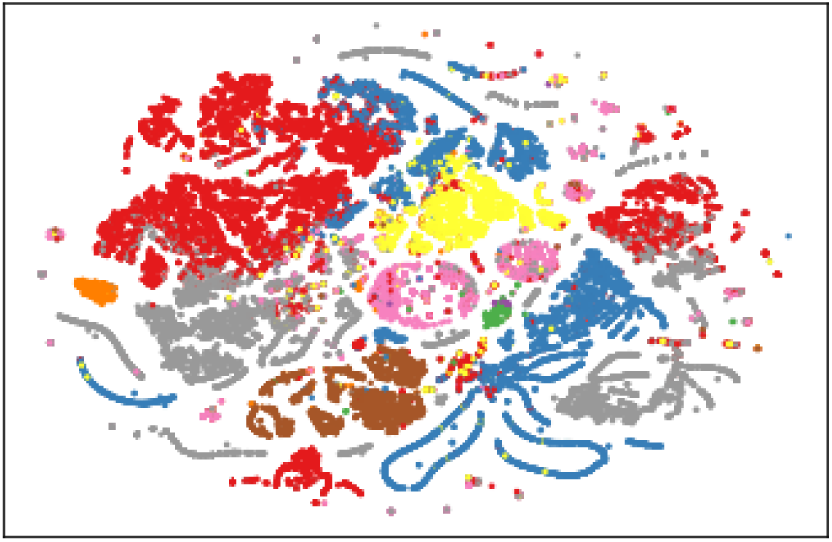
*t*-SNE Plot for Green Dataset using 11 features selected by CIFE (binary)

The error rates in 5-fold cross validation with random forests are summarized in Figures 7, 8, 10, and 9 (the corresponding figures for the nearest centroids classifier are Figures 13, 14, 15, and 16 at the end of the paper).

**Figure 7:**
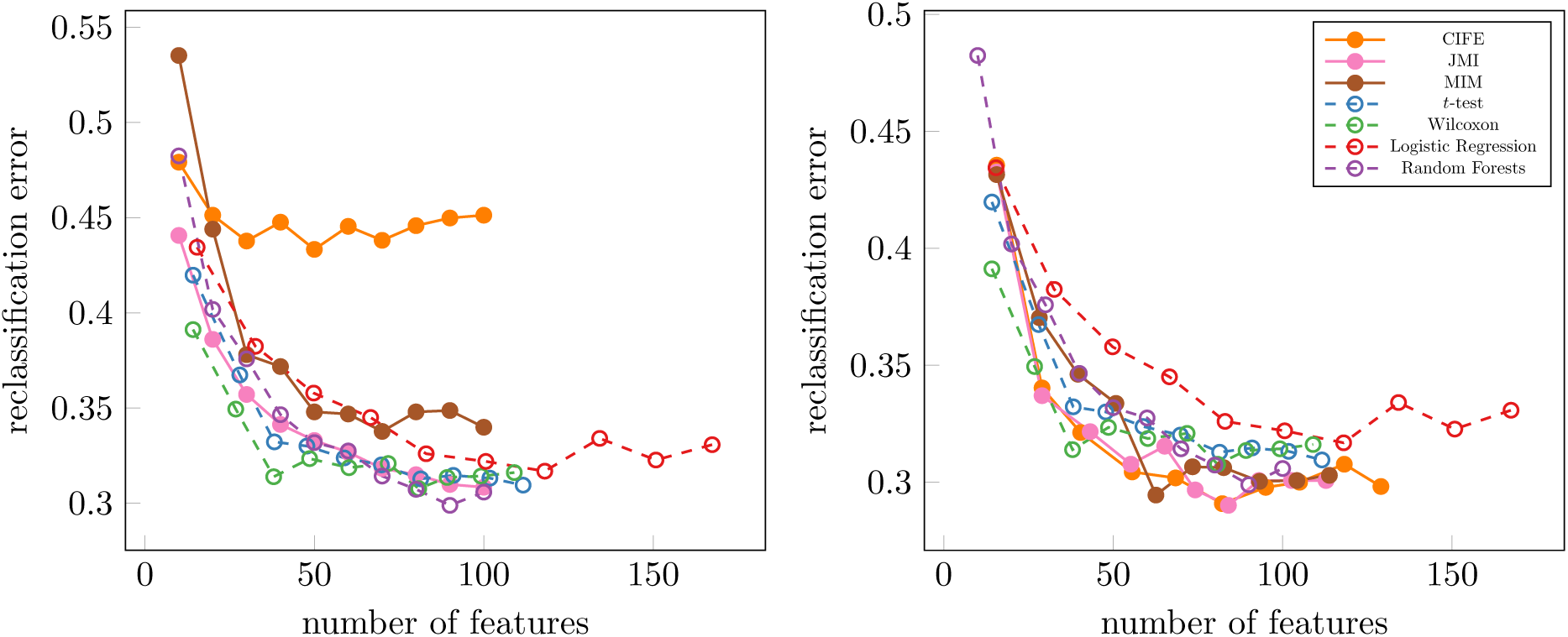
Classification Errors For Paul Dataset (left: multiclass MI methods, right: binary MI methods)

**Figure 8:**
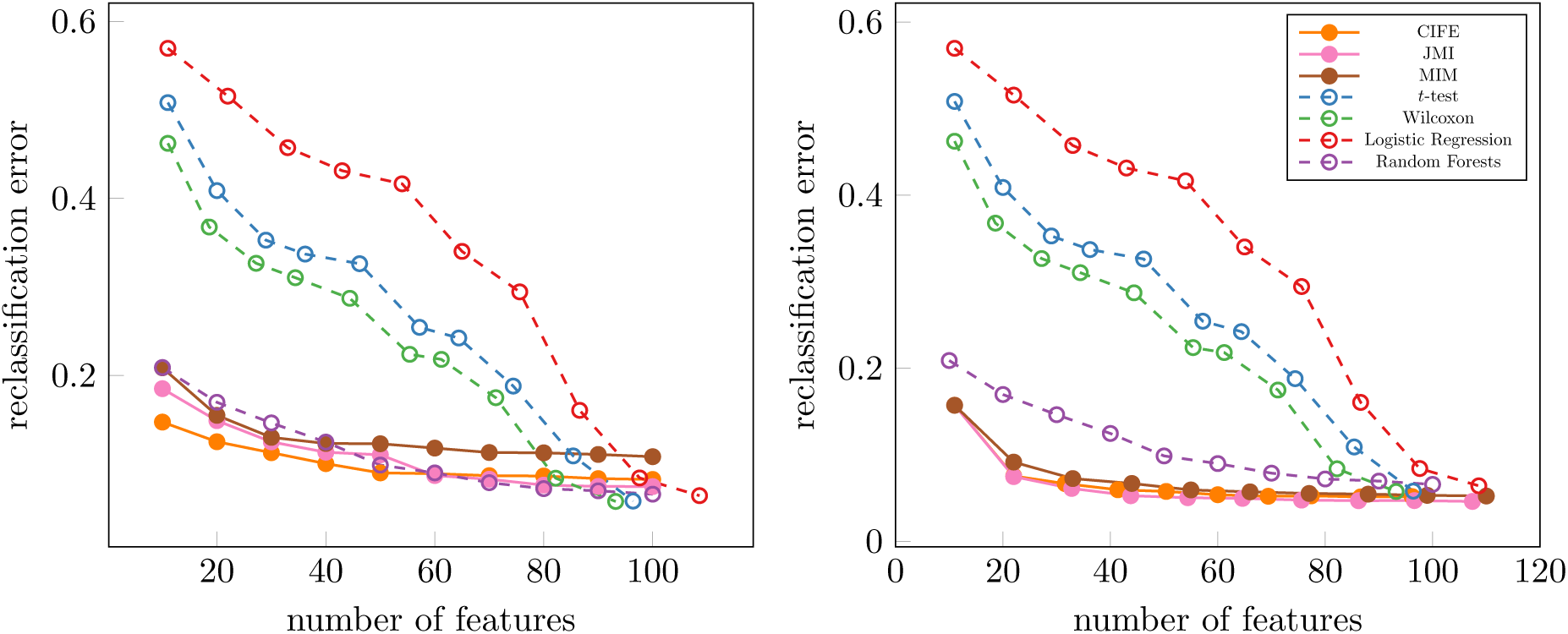
Classification Errors For Green Dataset (left: multiclass MI methods, right: binary MI methods)

**Figure 9:**
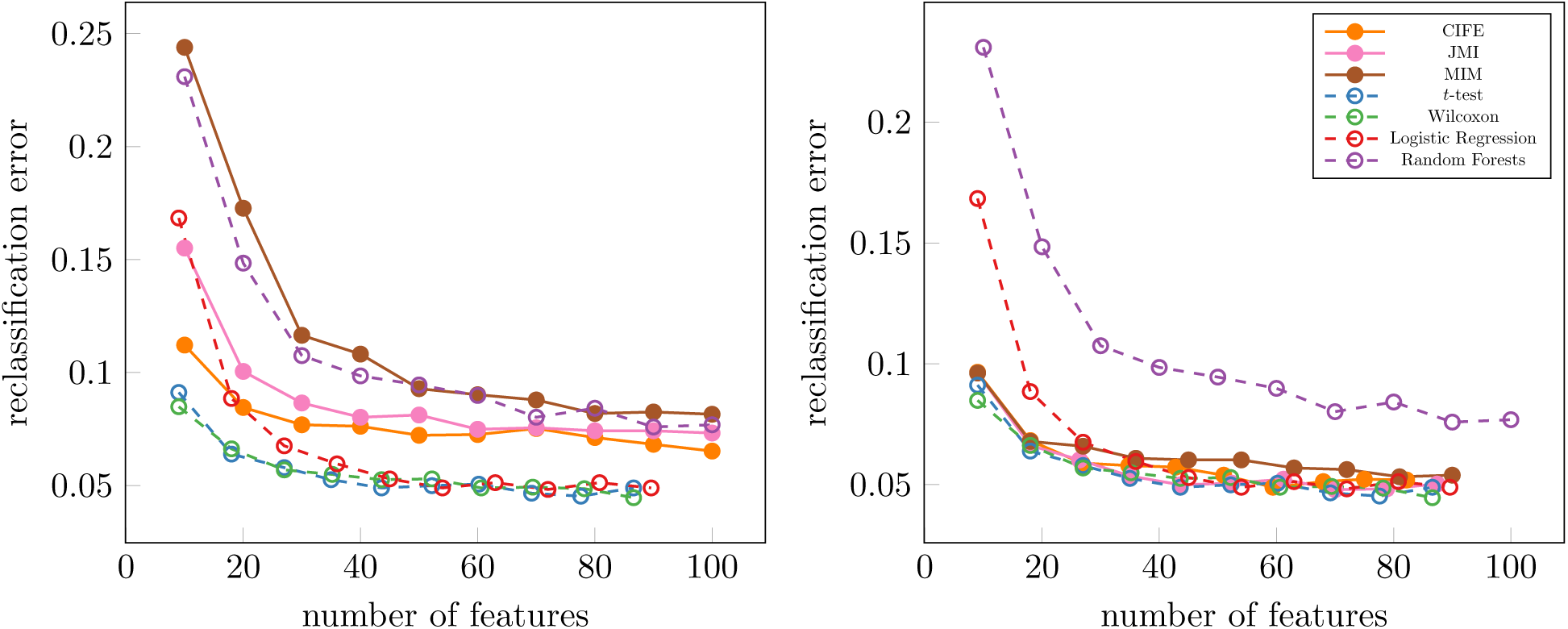
Classification Errors For Zeisel Dataset (left: multiclass MI methods, right: binary MI methods)

**Figure 10:**
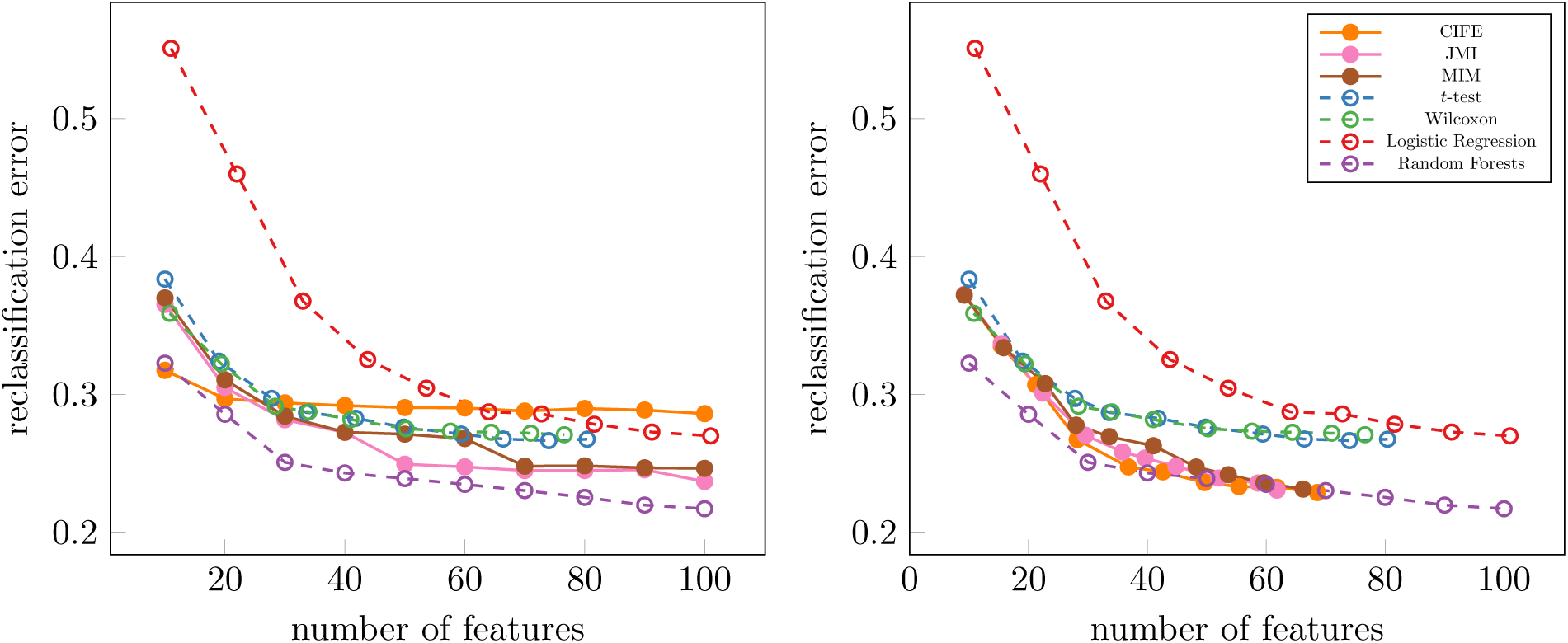
Classification Errors For Zheng Dataset (left: multiclass MI methods, right: binary MI methods)

**Figure 11:**
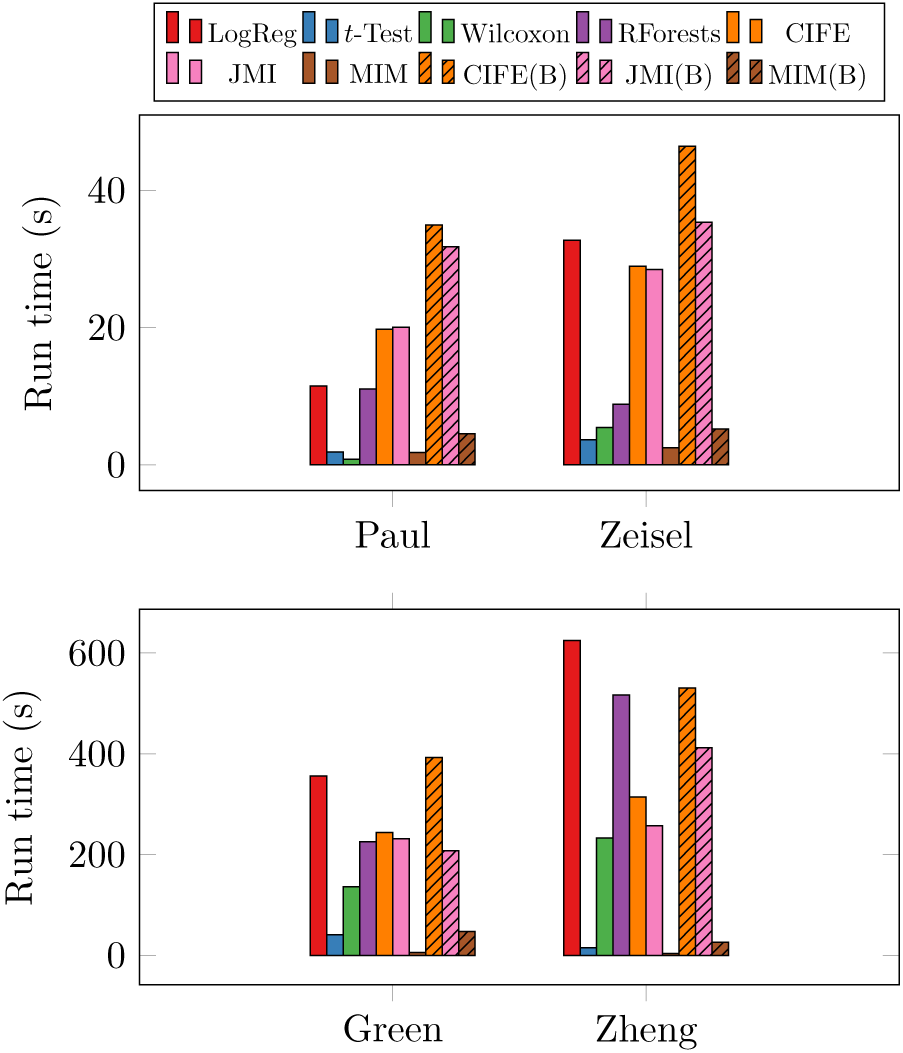
Runtimes for various algorithms

**Figure 12:**
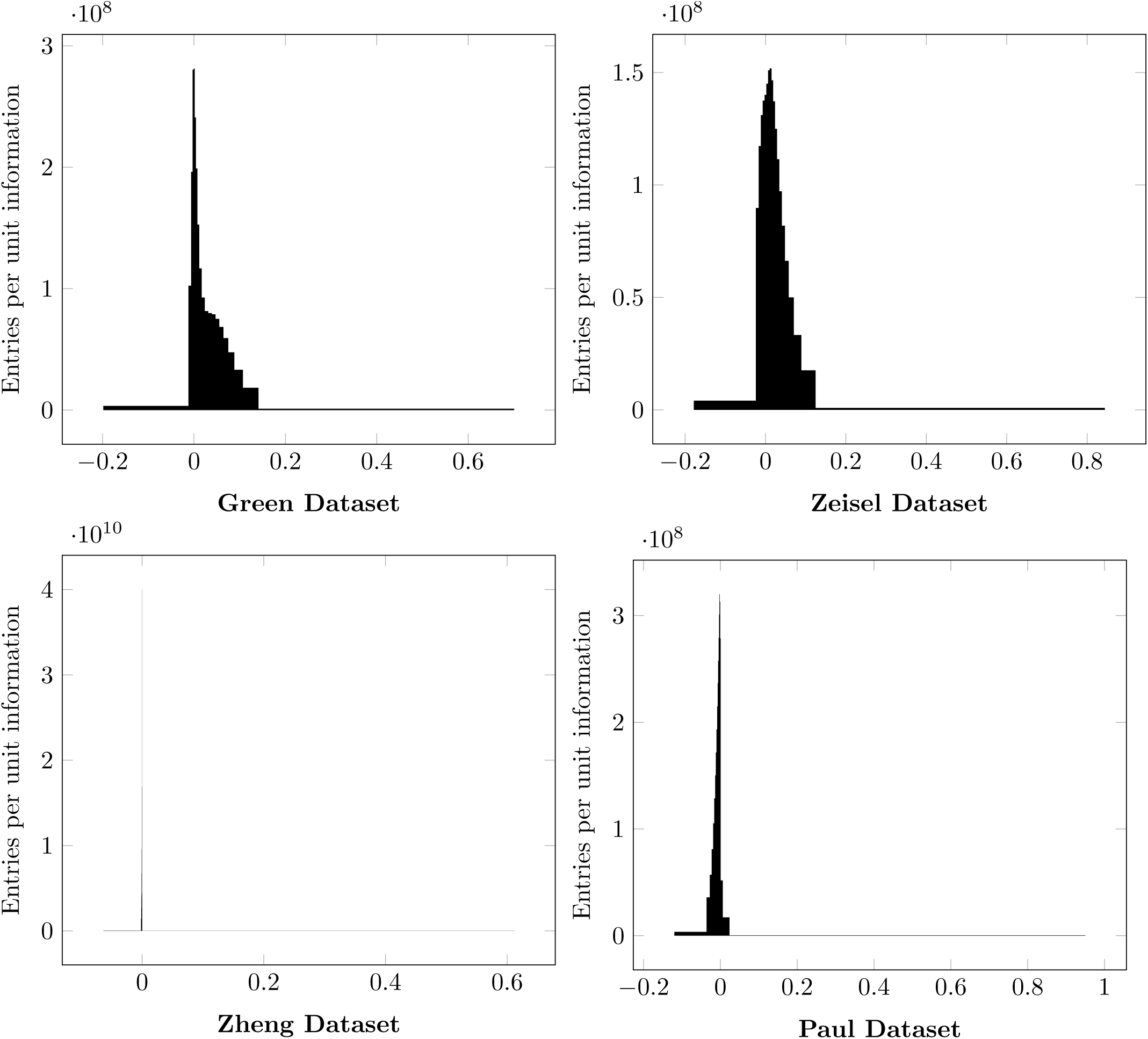
Histogram of *I*(*x*_*i*_; *x*_*j*_; *y*) values for various datasets.

**Figure 13:**
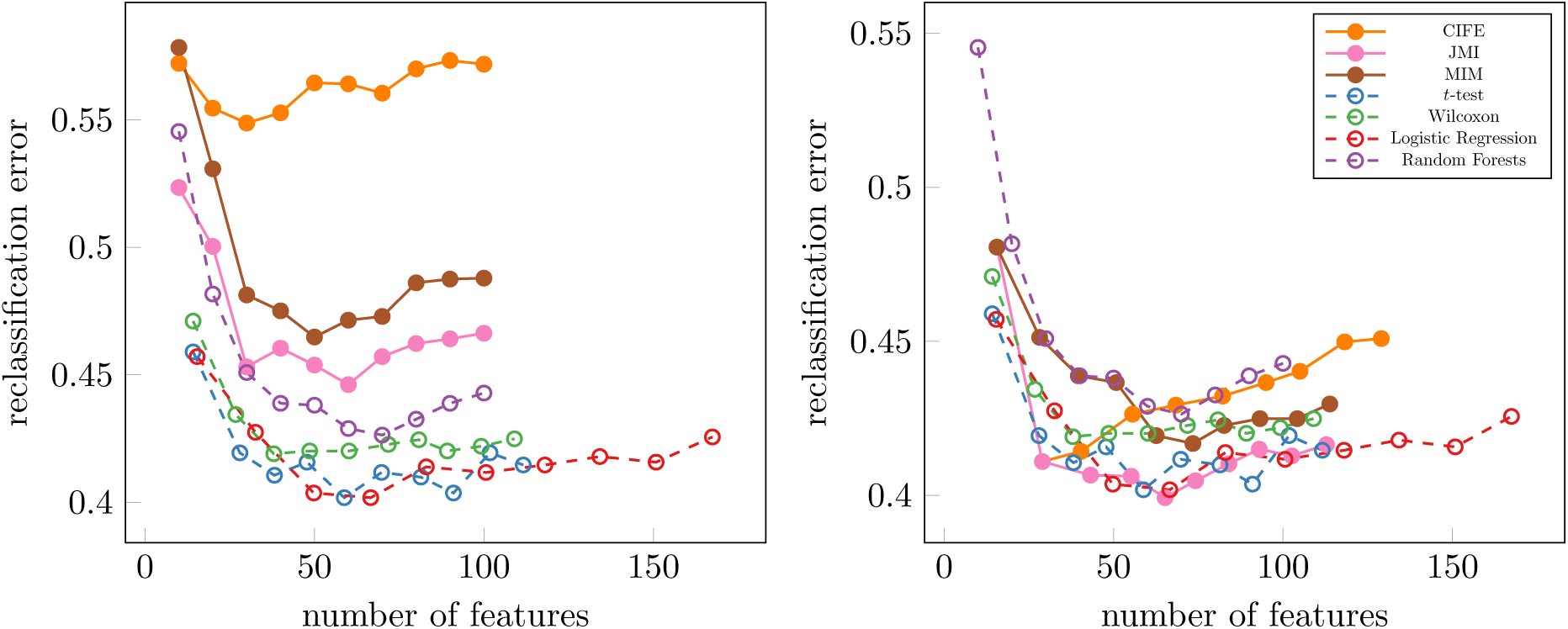
Classification Errors with Nearest Centroid Classifier For Paul Dataset (left: multiclass MI methods, right: binary MI methods)

**Figure 14:**
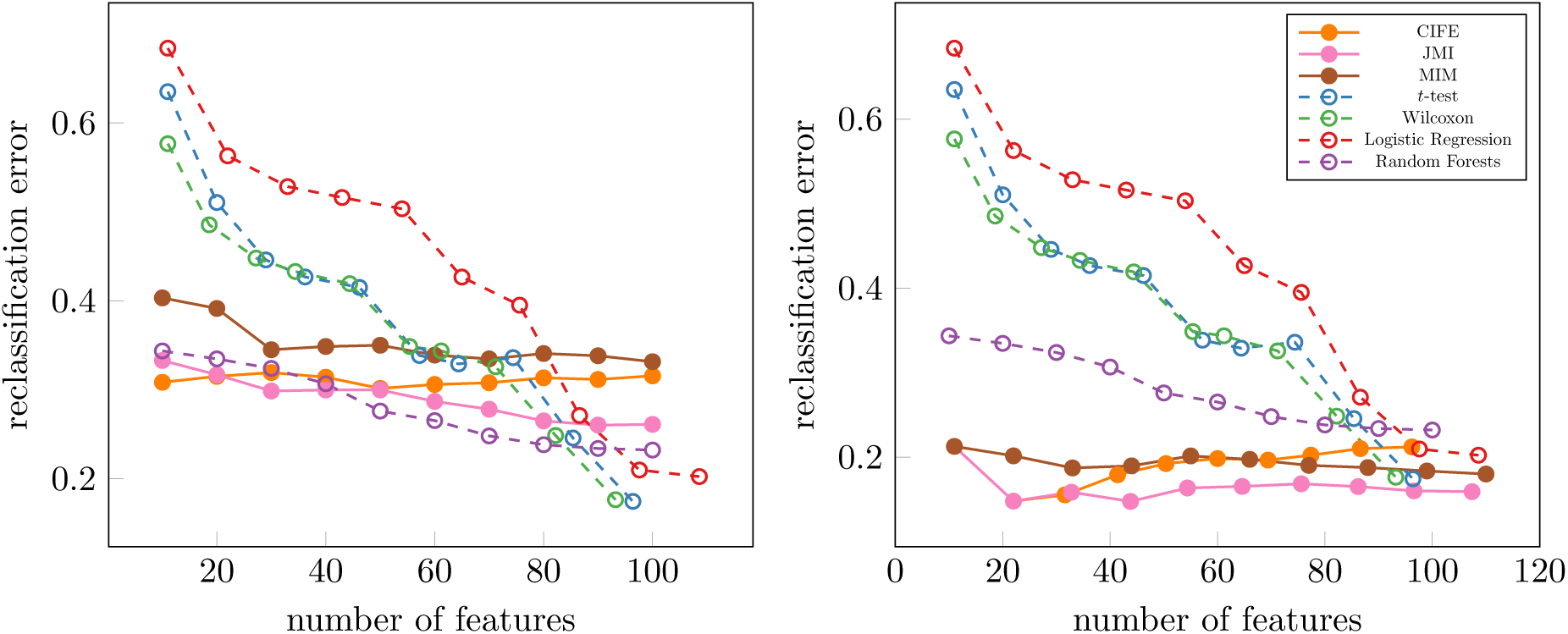
Classification Errors with Nearest Centroid Classifier For Green Dataset (left: multiclass MI methods, right: binary MI methods)

**Figure 15:**
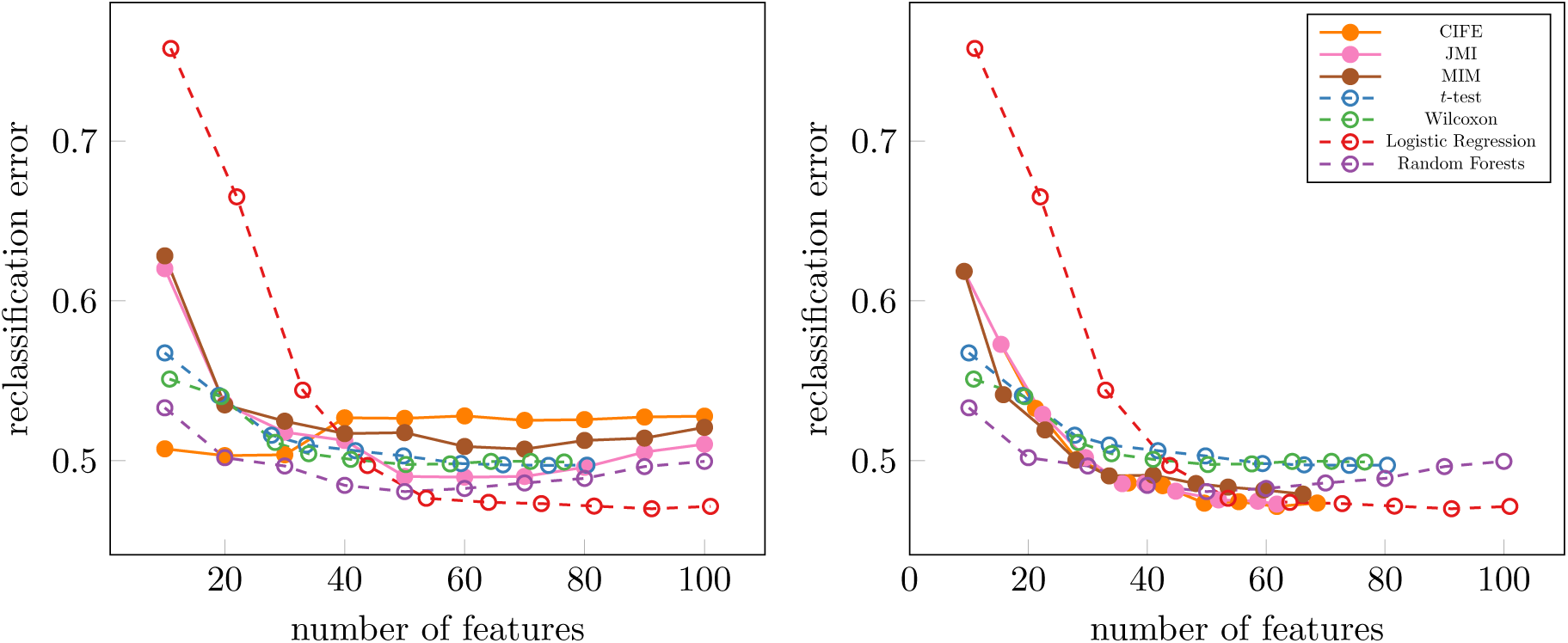
Classification Errors with Nearest Centroid Classifier For Zheng Dataset (left: multiclass MI methods, right: binary MI methods)

**Figure 16:**
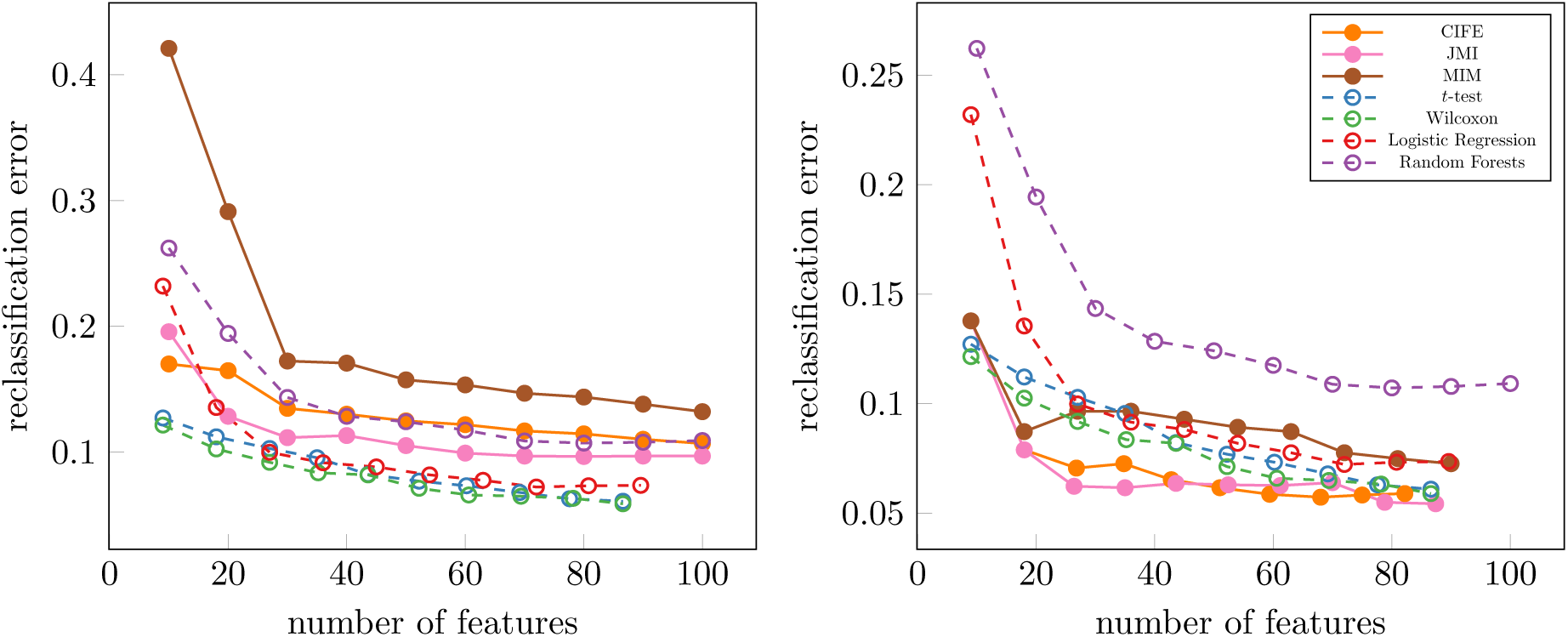
Classification Errors with Nearest Centroid Classifier For Zeisel Dataset (left: multiclass MI methods, right: binary MI methods)

In addition to error rates, which suggest how well the algorithms select features that explain class labels, we are also interested in knowing the extent to which these algorithm select the same features. We summarize these intersections in Tables 1, 2, 3, and 4. Because binary methods merge lists of features, we cannot ensure that all feature sets are of equal size. Therefore, not all sets are exactly of size 100 features, but they are the smallest possible sets of size ≥ 100.

**Table 1:**
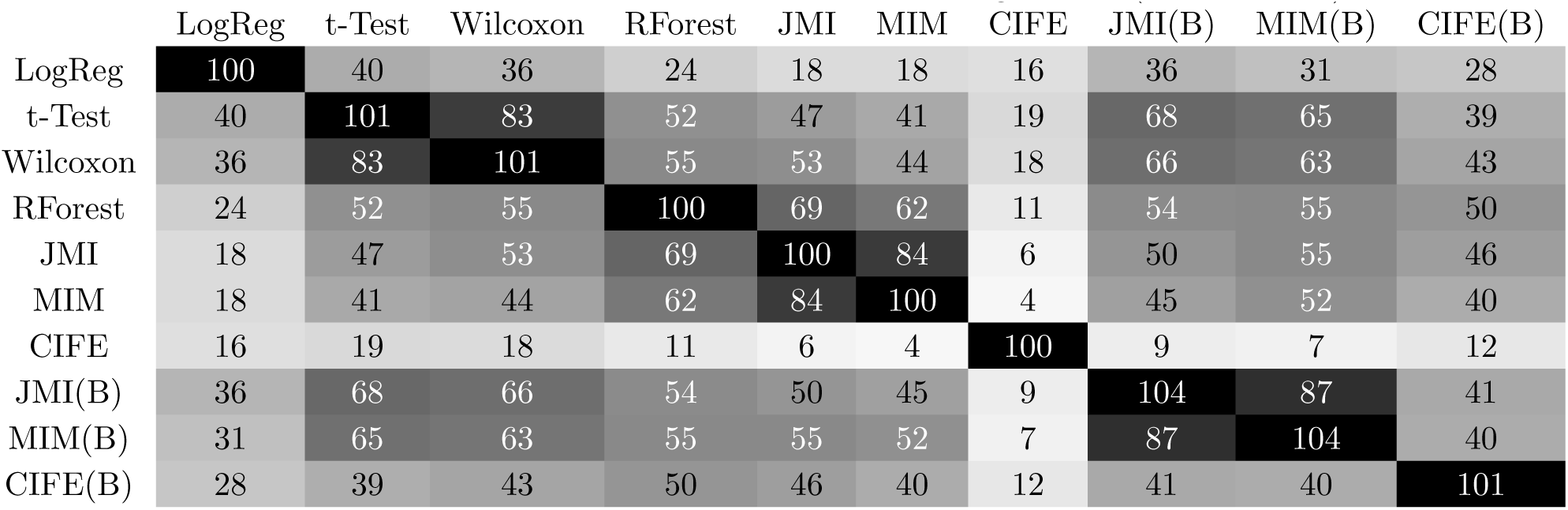
Number of Genes in Common between algorithms (Paul Dataset)

**Table 2:** Number of Genes in Common between algorithms (Green Dataset)

|  | LogReg | t-Test | Wilcoxon | RForest | JMI | MIM | CIFE | JMI(B) | MIM(B) | CIFE(B) |
| --- | --- | --- | --- | --- | --- | --- | --- | --- | --- | --- |
| LogReg | 108 | 38 | 32 | 8 | 9 | 7 | 3 | 49 | 45 | 20 |
| t-Test | 38 | 106 | 89 | 28 | 24 | 15 | 6 | 55 | 47 | 27 |
| Wilcoxon | 32 | 89 | 103 | 27 | 28 | 17 | 11 | 49 | 41 | 27 |
| RForest | 8 | 28 | 27 | 100 | 45 | 34 | 3 | 25 | 19 | 6 |
| JMI | 9 | 24 | 28 | 45 | 100 | 73 | 8 | 21 | 16 | 9 |
| MIM | 7 | 15 | 17 | 34 | 73 | 100 | 3 | 16 | 12 | 9 |
| CIFE | 3 | 6 | 11 | 3 | 8 | 3 | 100 | 4 | 1 | 18 |
| JMI(B) | 49 | 55 | 49 | 25 | 21 | 16 | 4 | 108 | 79 | 27 |
| MIM(B) | 45 | 47 | 41 | 19 | 16 | 12 | 1 | 79 | 100 | 23 |
| CIFE(B) | 20 | 27 | 27 | 6 | 9 | 9 | 18 | 27 | 23 | 109 |

**Table 3:**
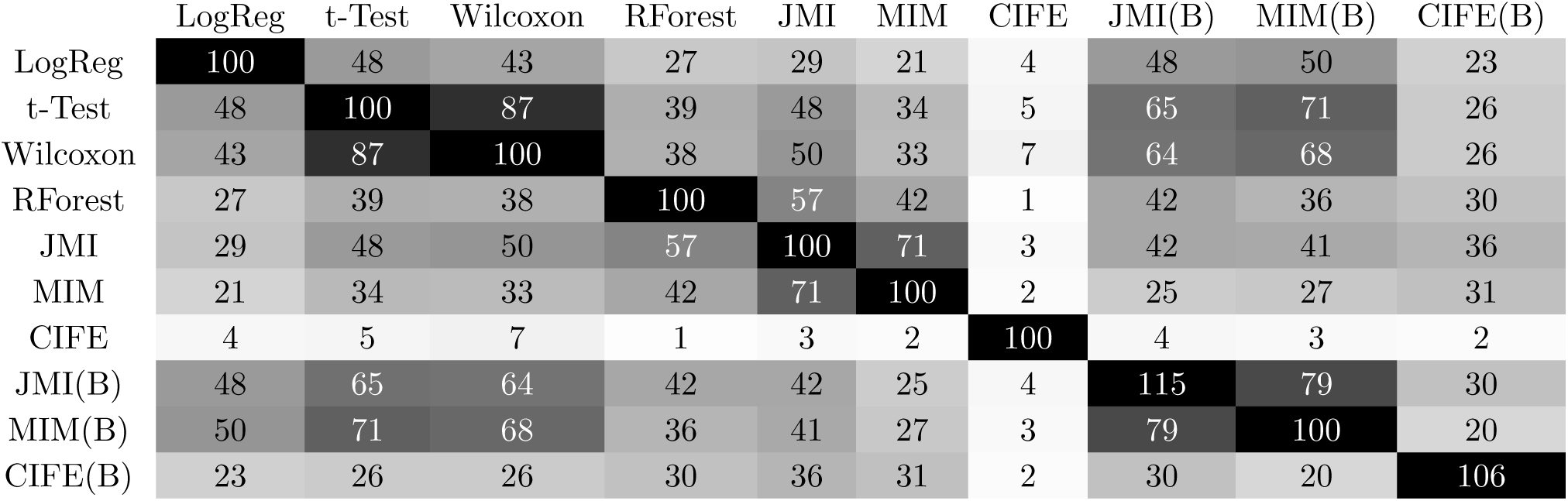
Number of Genes in Common between algorithms (Zeisel Dataset)

**Table 4:** Number of Genes in Common between algorithms (Zheng Dataset)

|  | LogReg | t-Test | Wilcoxon | RForest | JMI | MIM | CIFE | JMI(B) | MIM(B) | CIFE(B) |
| --- | --- | --- | --- | --- | --- | --- | --- | --- | --- | --- |
| LogReg | 102 | 31 | 29 | 19 | 19 | 18 | 2 | 33 | 31 | 19 |
| t-Test | 31 | 110 | 94 | 52 | 60 | 59 | 14 | 61 | 64 | 34 |
| Wilcoxon | 29 | 94 | 108 | 53 | 62 | 62 | 15 | 57 | 60 | 36 |
| RForest | 19 | 52 | 53 | 100 | 79 | 77 | 20 | 53 | 51 | 51 |
| JMI | 19 | 60 | 62 | 79 | 100 | 94 | 16 | 58 | 58 | 50 |
| MIM | 18 | 59 | 62 | 77 | 94 | 100 | 14 | 55 | 56 | 49 |
| CIFE | 2 | 14 | 15 | 20 | 16 | 14 | 100 | 14 | 9 | 28 |
| JMI(B) | 33 | 61 | 57 | 53 | 58 | 55 | 14 | 101 | 81 | 45 |
| MIM(B) | 31 | 64 | 60 | 51 | 58 | 56 | 9 | 81 | 101 | 37 |
| CIFE(B) | 19 | 34 | 36 | 51 | 50 | 49 | 28 | 45 | 37 | 100 |

In order to compare the run times for the various algorithms, we timed individual runs of each algorithm on each dataset (without any cross-validation). The recorded times for bivariate methods (the multiclass and binary variants of both CIFE and JMI) include the run time for the initial univariate selection of 5000 features with MIM.

## 3 Discussion

On the datasets we have evaluated, the mutual information based methods seem to consistently outperform differential expression methods when run as binary methods. However, when run as multiclass methods, mutual information based methods perform better than differential expression methods only on some datasets, e.g., the Green dataset (Figure 8) and the Zheng dataset (Figure 10), while differential expression methods far outperformed multiclass methods in other datasets, e.g., the Paul dataset (Figure 7). It is worth noting that it is only on the Zheng dataset that selecting features based on Random Forests’ feature importances is significantly better than other methods. We would also add that Figures 1, 2, 3, 4, and 5 demonstrate the strength of multiclass information theoretic methods on the Green dataset, but as we see by comparing Figure 8 to Figure 7, we don’t expect this to be the case on all datasets.

Tables 1, 2, 4, and 3 suggest that many binary methods choose many of the same features. The notable exceptions are logistic regression and the binary CIFE method, both of which have moderate overlaps with other methods, but still select numerous features that other methods do not. The multiclass CIFE algorithm has, by far, the most distinct feature sets. This proves to be both a source of strength and weakness as the classification errors associated to these feature sets are sometimes very poor (e.g., in the Paul dataset, Figure 7, and, to a much milder extent, the Zheng dataset, Figure 10) but sometimes very competitive (e.g., in the Green dataset, Figure 8, and in the Zeisel dataset, Figure 9). This suggests that in some cases, CIFE can be used to discover novel markers genes for cell types and/or cell states in scRNA-seq.

The run times for information-theoretic algorithms, as demonstrated by Figure 11, are comparable to their differential expression counterparts. Specifically, the univariate MIM is almost always the fastest and the bivariate CIFE and JMI have similar run times to logistic regression.

Finally, we consider the assumptions we made to give our theoretical guarantees in the Guarantees for Greedy Search section and examine how close these datasets are to satisfying those assumptions. One of the most significant assumption we made (e.g., in Corollary 14) is that *I*(*x*_*i*_; *x*_*j*_; *y*) ≥ 0 for all *x*_*i*_, *x*_*j*_ ∈*𝒳*. In Figure 12, we see that although the hypothesis don’t hold perfectly on any dataset, they hold best on the Green dataset (where CIFE seems to perform best, see Figure 8) and worst on the Paul dataset (where CIFE seems to perform the worst, see Figure 7).

## 4 Conclusion

The opportunity presented by single cell RNA sequencing also poses challenges in its analysis. While traditional differential expression reveal meaningful information about the data, we believe that a multiplicity of feature selection techniques allows researchers to discover novel marker genes. Specifically, we believe that information-theoretic feature selection algorithms can detect marker genes that may otherwise be obscured by interactions with other genes or distributional assumptions of statistical tests.

We do not expect (in fact, we believe it is unrealistic to hope) that a single marker selection technique is appropriate for all purposes and datasets. The algorithms in this paper (especially CIFE) have proven to select sets of genes that are very distinct from those selected by traditional differential expression methods. In most cases the reclassification accuracy when restricting datasets to the chosen genes was comparable between information-theoretic algorithms and traditional differential expression algorithms and in some cases information-theoretic algorithms performed considerably better.

The reader will likely have noted that there are numerous possible combinations of configurations and versions of these algorithms to choose from. We attempt to summarize these options in the form of suggestions below.

- MIM is a univariate algorithm. Because it does not consider interactions between genes, it yields somewhat similar results to traditional differential expression algorithms, especially when run as a binary method rather than a multiclass method. As a multiclass method, it is particularly susceptible to selecting features with a strong signal without any consideration of how informative those features are together or what class labels they are informative about. Therefore, the output of MIM is more meaningful and easier to interpret when it is run as a binary method.
- CIFE is a bivariate algorithm that aggressively penalizes redundancy. When used as a multiclass method, it is particularly strong at identifying a small set of informative genes (in our examples, CIFE often far outperformed other algorithms when selecting 10 to 30 genes). CIFE’s ability to consider interactions allows it to pick genes that may not be as informative in isolation but are informative in the context of other selected genes. However, when selecting a large number of genes, the interactions (particularly redundancies) overwhelm the algorithm and it eventually picks very uninformative features. This is especially true when individual genes have information about numerous class labels (for example, when there are many class labels in the dataset; see Example 5). If we need a larger set of genes or want marker genes for specific class labels, we recommend using CIFE as a binary feature selection method. Running the algorithm as a binary method ensures that the number of features chosen per class label remains relatively low.
- JMI is a bivariate algorithm that acts as an intermediate between CIFE and MIM. As the algorithm selects more features, it dampens the impact of interactions resulting, eventually, in a convergence in behavior between JMI and MIM. As a consequence of this damping, it is unlikely to unearth informative features hidden behind interactions as CIFE can nor does its output reflect the strength of the signal of features in isolation. The damping in JMI is a heuristic that is not justified theoretically and should be used cautiously. However, empirical results show that its performance is consistently decent and avoids the occasional extreme poor performance of CIFE.

In all of these cases, we strongly recommend using the recursive quantile transform we describe under Binning Technique to discretize datasets.

Used in conjunction with traditional differential expression methods, the information-theoretic algorithms in this paper have the potential to discover novel marker genes that reveal information that otherwise be lost on differential expression algorithms that prioritize signal strength of genes over new information. Future directions include more sophisticated analysis and benchmarking of unsupervised feature selection methods for single cell RNA sequencing (see supplementary materials for examples).

## 5 Methods

Our benchmarking process relied heavily on scanpy, a widely used package for single cell analysis in Python, and PicturedRocks, a software package written by the first and last authors of this paper. PicturedRocks is a Python package that can perform information-theoretic feature selection and also contains tools to benchmark feature selection methods via *k*-fold cross-validation. PicturedRocks includes a interactive user interface for feature selection that can allows researchers to incorporate domain knowledge in the process of feature selection. All the code to reproduce our benchmarking study can also be found on GitHub.

There are numerous possible pitfalls when evaluating the ability of a feature selection method to select informative features. We refer the reader to the excellent section on “The Wrong and Right Way to do Cross-validation” in [18, Section 7.10.2]. In particular, we highlight the following:

- Cross-validation is effective at estimating the generalization error rather than the test error, which is susceptible to artificially being suppressed due to overfitting.
- It is extremely important to ensure that at no point does any data “leak” between folds. For example, as explained in [18], it is not fair to evaluate a feature selection method by first selecting features on the entire dataset and then report the cross-validation error. Even though the classifier used to predict labels for each fold was not trained on that fold, the feature selection algorithm had access to data from all folds.

In order to avoid such pitfalls, we performed our benchmarking in the following manner:

1. Split the dataset into 5 folds
2. For each fold *k* = 1, *…*, 5:
  a. use all samples not in fold *k* to select *n* features; repeat for each feature selection algorithm and store results separately *Remark 1:* For binary feature selection methods (such as *t*-test, Wilcoxon Rank Sum, Logistic Regression, and when any of the information-theoretic methods are forcibly run as binary methods), this means running the algorithm for each class label and combining the top ⌈*n/c*⌉ features from each class label, where *c* is the number of classes. Remove duplicates. Keep track of the number of features selected, since this may not be exactly *n*. *Remark 2:* the bivariate methods (CIFE and JMI) can slow down considerably in situations when we are selecting many features and there are many candidate features (specifically there is a *O*(*np*) term in the time complexity of these methods). To avoid these prohibitively slow run times, we use a univariate method (MIM) to select 5000 features and then use the bivariate method to select among those 5000 features
  b. train a Random Forests classifier and a Nearest Centroids classifier on all samples not in fold *k* with only the *n* (or possibly a few more) features selected; repeat for each feature selection algorithm
  c. use both classifiers to predict labels for samples in fold *k* and store the predictions of each classifier separately; repeat for each feature selection algorithm
3. for each feature selection algorithm and classifier combination:
  a. compute the error rate of the predictions as #(wrongly classified)*/*#(samples).
  b. for binary methods where the number of features may not be exactly *n*, record the number of features used as the average number of features used for each fold

For *t*-test, Wilcoxon Rank Sum, and Logistic Regression feature selection methods we used the standard implementation in scanpy. As part of the feature selection subroutines, we perform basic preprocessing (as this happens on a copy of the data, it does not affect the data used for classification): normalize per cell followed by log1p. These methods scale the counts matrix so that each cell’s counts sum to the same number (this is usually chosen to the median count prior to scaling) and then apply the *x* ↦ log(1 + *x*) transformation element-wise. For the information-theoretic methods, we use the recursive quantile transformation (with 5 bins) before running the methods, see Binning Technique in the supplementary material of this paper.

At the classification step, we used the Random Forests classifier from the Scikit-learn package[16] with 100 estimators. As a preprocessing step, we scaled the data so that the sum of counts for each cells was 1000 (this is arbitrary but had to be chosen in order to standardize the preprocessing between training and testing) and then applied the *x* ↦ log(1 + *x*) transformation. For the Nearest Centroids classifier, we applied the *x* ↦ log(1 + *x*) transformation, projected onto the top 30 principal components of the training data (as euclidean distances become less meaningful in high dimensions), and the mapped each training point to the class with the nearest centroid in the training data.

When generating Figure 11 and Tables 1, 2, 3, 4, we performed the preprocessing for feature selection algorithms listed above and ran the algorithms on the entire dataset rather than use cross validation as these figures are not meant to estimate generalizability. Figures 1, 2, 3, 4, 5, and 6 were also generated with these feature sets using the *t*-SNE implementation in Scikit-Learn [16].

### 5.1 Introduction to Information Theoretic Methods

The remainder of this section introduces the basic concepts behind information theoretic feature selection methods. In addition to benchmarking these methods, we have performed theoretical analyses of these algorithms and present our findings in detail in the supplementary material.

Recall that we seek to find an small subset of genes that is most informative about the class labels. In the language of information theory, we can frame this as the problem finding a small subset *S* of the variables 𝒳 (the set of all genes) such that *I*(*S*; *y*), the mutual information between a set of features *S* and target labels *y*, is maximized. This warrants some definitions before we proceed.

#### Definitions, Notation, and Abuses Thereof

We begin with the definition of mutual information. The mutual information between random variables *U* and *V* can be thought of as the number of bits of information knowing *U* tells us about *V*. For example, if *U* is a outcome of a fair coin toss (i.e., *U* takes values in {*H, T*} with equal probability) and *V* is the opposite of *U* (i.e., *V* is *T* if *U* is *H* and vice-versa), then their mutual information is 1 bit. However, if *V* is a “noisy” opposite of *U* so that *V* is the opposite of *U* with probability 0.99 and is equal to *U* with probability 0.01 then *I*(*U*; *V*) should be just under 1 (in fact it is approximately 0.92).

##### Definition

If *U* and *V* are discrete random variables, then the mutual information *I*(*U*; *V*) between *U* and *V* is

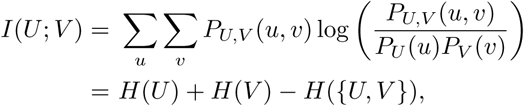

where

- *P*_*X*_ is the probability mass function of *X*, a random variable, in particular *P*_*U,V*_ (*u, v*) = *P* (*U* = *u* and *V* = *v*),
- *H*(·) is the Shannon entropy of a discrete random variable, and
- the above logarithms are with base 2, by convention.

We will only consider the case when *U* and *V* are discrete random variables (in practice, such algorithms are used after data has been discretized, although continuous analogues that use density estimators exist, but are computationally harder).

By abuse of notation and in line with existing literature, we will often

- treat a set of random variables as an ensemble of random variables. That is,

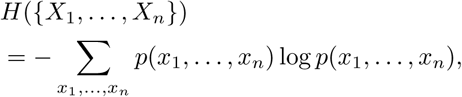
- drop the ∪ symbol and write *H*(*UV*) to mean *H*(*U* ∪ *V*), and
- treat a feature *x*_*i*_ as a singleton {*x*_*i*_}, so *H*(*Ux*_*i*_) = *H*(*U* ∪ {*x*_*i*_}).

#### Information-theoretic Feature Selection

We may now return to making the statement of our problem precise. We want to find *S* ⊂ 𝒳 such that *I*(*S*; *y*) is maximum over subsets of size *k* or less of 𝒳. Here *k* is a user-prescribed number of features to select. There are 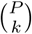 such subsets (for fixed *k*, this is *O*(*P*^*k*^)). Datasets often have many thousands of features from which we may seek to select on the order of a hundred features, which makes an exhaustive search computations intractable. It is already known that feature selection is NP-hard (see [8]), as are sub-modular maximization problems for coverage-type functions (see [19]) that our problem can be reduced to by constructions similar to those discussed under Analysis of Existing Methods in this paper. Various greedy algorithms, usually of the form of Algorithm 1, have been studied in the literature. We give a summary of these algorithms in Table 5. These algorithms use different objective functions *J*: 𝒳→ ℝ, where *J* (*x*_*i*_) estimates the (relative) value of feature *x*_*i*_ given that *S* ⊂ 𝒳 has already been chosen.

**Table 5:**
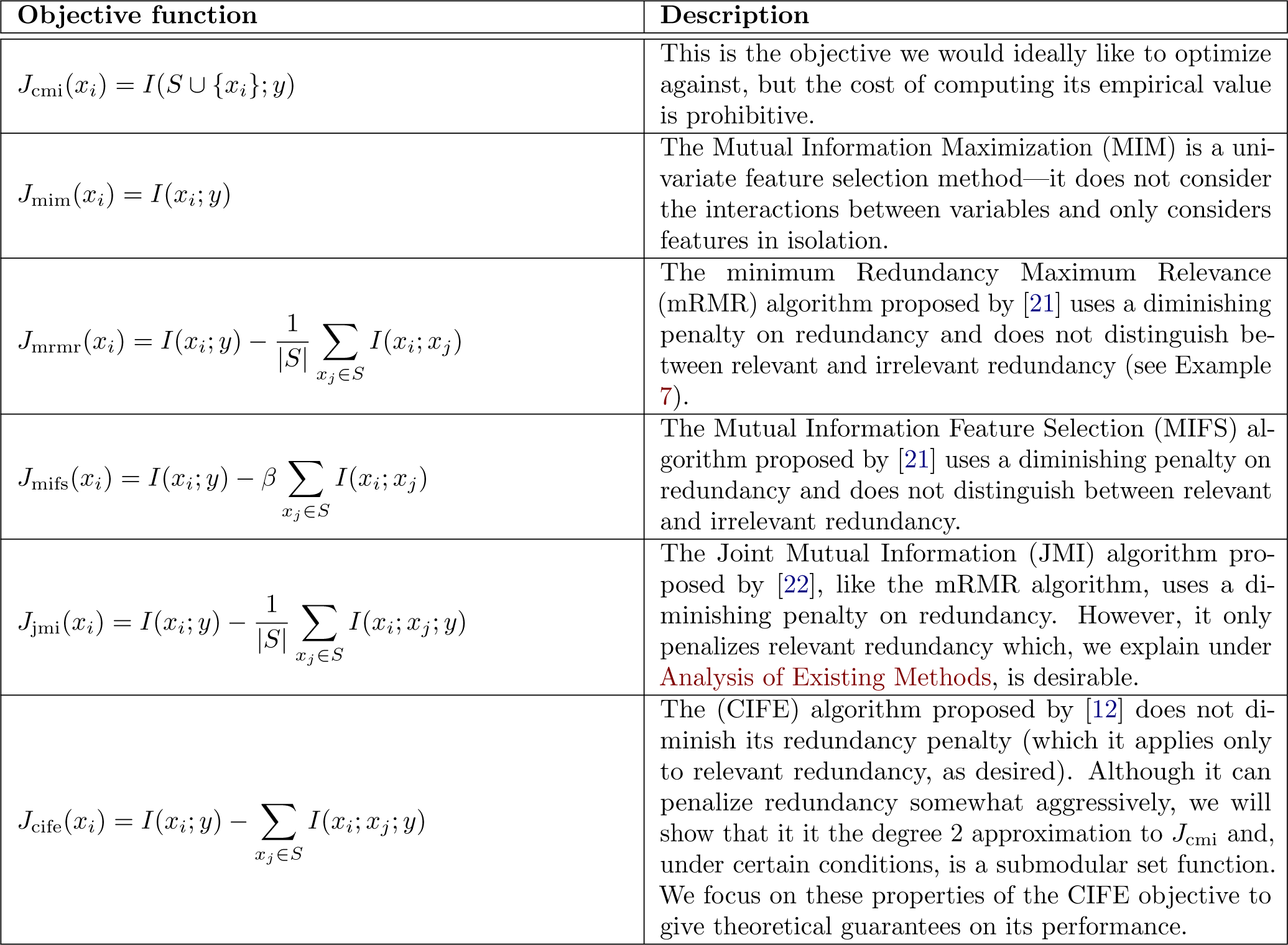
Summary of various objective functions that are used with Algorithm 1.

##### Algorithm 1 General Greedy Algorithm for Feature Selection with Objective *J*

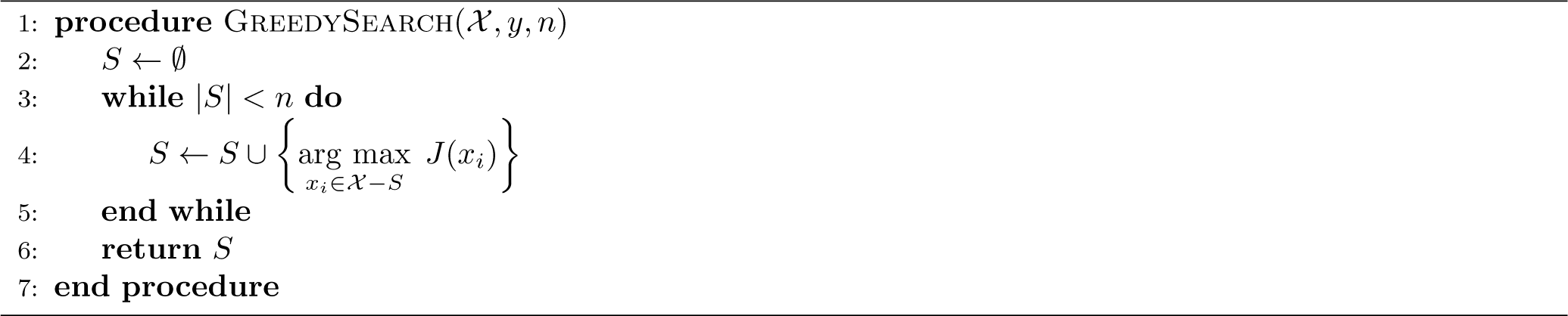

If we were to naïvely maximize *I*(*S*; *y*) via a greedy search, we would run Algorithm 1 with *J*_cmi_(*x*_*i*_) = *I*(*S*∪{*x*_*i*_}; *y*). Computing this requires estimating the joint probability distribution of *Sx*_*i*_*y* (which has *|S|*+ 2 dimensions). This is not only computationally intractable, but also impossible to estimate reasonably in the absence of a very large number of samples. Therefore, the objective function

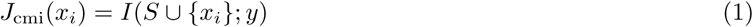

is difficult to estimate. We follow the terminology of Brown, et al. [20] and will refer to this as the conditional mutual information (CMI) objective because a greedy search with the conditional mutual information objective, *J* (*x*_*i*_) = *I*(*x*_*i*_; *y|S*), is equivalent to using *J*_*cmi*_ from (1). This is because *I*(*x*_*i*_; *y|S*) *– I*(*Sx*_*i*_; *y*) is constant with respect to *x*_*i*_.

In this work, we present a series expansion of *I*(*S* ∪ {*x*_*i*_}; *y*) in terms of multivariate mutual information through which we can derive degree *K* approximations to *I* (*S* ∪ {*x*_*i*_}; *y*). This expansion gives us insight into the strengths and weaknesses of various objective functions in the literature. We then focus on the degree 2 approximation, which corresponds to the CIFE feature selection algorithm [12], and give theoretical guarantees for its performance. We then evaluate the performance of these algorithms on publicly available scRNA-seq datasets and describe optimizations for sparse data that improve the computational efficiency of these algorithms considerably.

### 5.2 Previous work

Because of the computational challenges in empirically evaluating *J*_cmi_, many approximation algorithms have been proposed. Brown et al. [20] generalize a vast number algorithms in the literature as

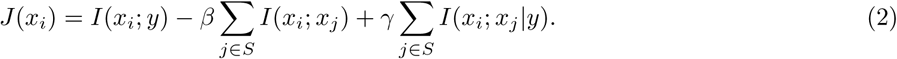

Specifically, the widely used mRMR algorithm in [21] has 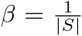 and *γ* = 0. The MIFS algorithm by [23] has *γ* = 0 and leaves *β* as a (constant) hyperparameter chosen from [0, 1]. The CIFE algorithm in [12] has *β* = *γ* = 1. The JMI algorithm ([22]) and IGFS^2^ algorithm ([24]) use 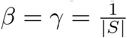. See Table 5 for a summary of the various algorithms described.

Brown et al. provide an interpretation of this class of objective functions in terms of three assumptions.

1. Given an unselected feature *x*_*k*_,
  a. the selected features are independent: 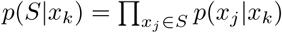 and
  b. the selected features are class-conditionally independent: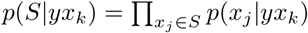.
2. Features are pairwise class-conditionally independent: *p*(*x*_*i*_*x*_*j*_*|y*) = *p*(*x*_*i*_*|y*)*p*(*x*_*j*_*|y*).
3. Features are pairwise independent: *p*(*x*_*i*_*x*_*j*_) = *p*(*x*_*i*_)*p*(*x*_*j*_).

### 5.3 Our Work

We attempt to address two sets of important questions about the information theoretic methods that we discuss in this paper:

- What are the differences between the various objective functions? Under what conditions does the CMI objective function (the ideal objective function that we would use if computing power and quantity of data were not restricted) reduce to the approximate objective functions such as MIM, mRMR, JMI, and CIFE? What explains the relative strengths and weaknesses of these objective functions?
- What guarantees can we make for the greedy search algorithm? How far from optimal are the outputs of the greedy search?

We address the first point by expanding the CMI objective function *J* (*x*) = *I*(*S* ∪ {*x*}, *y*), which considers all possible interactions, in terms of lower-order interactions under the Series Expansion of Mutual Information section. We show that truncating this series after the degree 2 terms gives us the CIFE objective. The reader may also find it helpful to read the example-based section on Analysis of Existing Methods which demonstrates where various objective functions fail.

The second set of question, on the effectiveness of the greedy search algorithm, we demonstrate in the section titled Guarantees for Greedy Search that under certain conditions, mutual information and its degree 2 approximation (i.e., CIFE) are submodular functions and existing literature on submodular functions to give theoretical guarantees for the performance of the greedy search algorithm for CIFE.

Under Practical Implementation Issues, we explain various considerations that the practitioner will find useful. This includes a section our approach to preprocessing datasets for information-theoretic feature selection methods as these methods require the counts to be in discrete bins, as well as a section on trade offs between running information-theoretic methods as mutli-class and binary methods.

Finally, we note that we have implemented fast versions of these algorithms in the PicturedRocks Python package and we explain some of the optimizations that we made under Efficient Algorithms.

## Availability of data and material

Our package PicturedRocks is available on GitHub and the Python Package Index (pip install picturedrocks) and installs scanpy as a dependency. The Paul dataset [13] is available on the NCBI Gene Expression Omnibus (GEO: GSE72857) and also ships with scanpy. The Green dataset [14] is available on the NCBI Gene Expression Omnibus (GEO: GSE112393). The Zheng dataset ships with scanpy and is available on 10x Genomics website (https://support.10xgenomics.com/single-cell-gene-expression/datasets). The Zeisel dataset [15] is available on the Linnarsson Lab webpage (http://linnarssonlab.org/cortex/). Code for all the benchmarking experiments performed in this paper is available at https://github.com/umangv/picturedrocksbenchmarks.

## Competing interests

The authors declare that they have no competing interests.

## Author’s contributions

UV developed PicturedRocks and wrote the code for benchmarking and analyzed the data with assistance from JC and ACG. UV developed the theoretical analysis with assistance from ACG. All authors contributed to the writing of the manuscript and approve the final manuscript.

## Funding

UV and ACG are supported in part by the Michigan Institute for Data Science, and the Chan Zuckerberg Initiative. JC is supported by grants from the National Institute of Environmental Health Sciences (R01 ES028802) and the Michigan Institute for Data Science.

## Acknowledgements

The authors acknowledge the help of Jun Z. Li and Alexander Vargo for helpful discussions.

## Figures

### A Series Expansion of Mutual Information

In the Ignoring Higher-degree Interactions Between Features section, we give examples of *shared redundancy* and *synergy*. These interactions between features is explained by multivariate mutual information. Intuitively, we want to measure how much information a set of random variables share simultaneously. If the one bit of information that *x*_3_ shares with *x*_1_ is the same as the one bit of information that *x*_3_ shares with *x*_2_, then we want *I*(*x*_1_; *x*_2_; *x*_3_) = 1, and if *x*_3_ shares different bits of information with *x*_1_ and *x*_2_, then *I*(*x*_1_; *x*_2_; *x*_3_) = 0.

This suggests a generalization of mutual information that was studied in [25]. We let *I*_1_(*x*) = *H*(*x*) to be the Shannon entropy of *x* (i.e., the self-information). Define the multivariate mutual information *I*_*n*_(*x*_1_; *…*; *x*_*n*_) recursively as *I*_*n*_(*x*_1_; *…*; *x*_*n*_) = *I*_*n–*1_(*x*_1_; *…*; *x*_*n–*1_) *– I*_*n–*1_(*x*_1_; *…*; *x*_*n–*1_*|x*_*n*_), where *I*_*n–*1_(*x*_1_; *…*; *x*_*n–*1_*|x*_*n*_) is the information shared between *x*_1_, *…, x*_*n–*1_ given we know the value of *x*_*n*_. If *S* = {*x*_1_, *…, x*_*n*_}, we denote by *I*_*n*_(*S*) the multivariate mutual information *I*_*n*_(*x*_1_; *x*_2_; *…*; *x*_*n*_). Note the subtle, but crucial, difference in notation: *I*_*n*_(*S*) is the information simultaneously shared by the *n* variables in *S* (intuitively, the intersection of information), while *I*(*S*) is the self-information of the ensemble of random variables of *S* (intuitively, the union of the information). The following equality (`a la Principle of Inclusion and Exclusion) expresses the multivariate mutual information *I*_*n*_(*S*) in terms of lower degree information terms.

#### Proposition 1

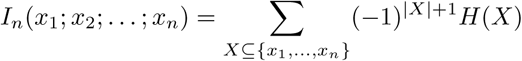

*Proof.* We prove the equality above by induction. For *n* = 2, this follows from the definition of mutual information. For *n >* 2, we have

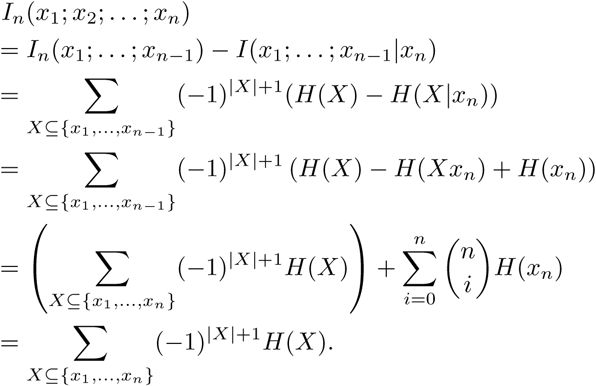

□

#### Corollary 2

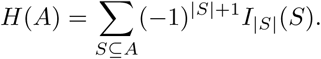

*Proof.* This follows from a version of the principle of inclusion and exclusion, see [26, p. 1049 Theorem 12.1]. □

We now give a series expansion of the mutual information *I*({*x*_1_, *…, x*_*n*_}, *y*) that suggests approximations to *I*(*x*_1_, *…, x*_*n*_, *y*) using lower degree terms (i.e., without needing to estimate the joint probability distribution of a large set of variables).

**Lemma 3.**

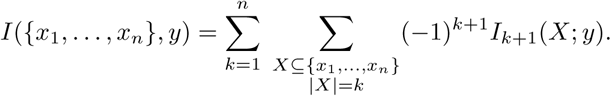

*Proof.*

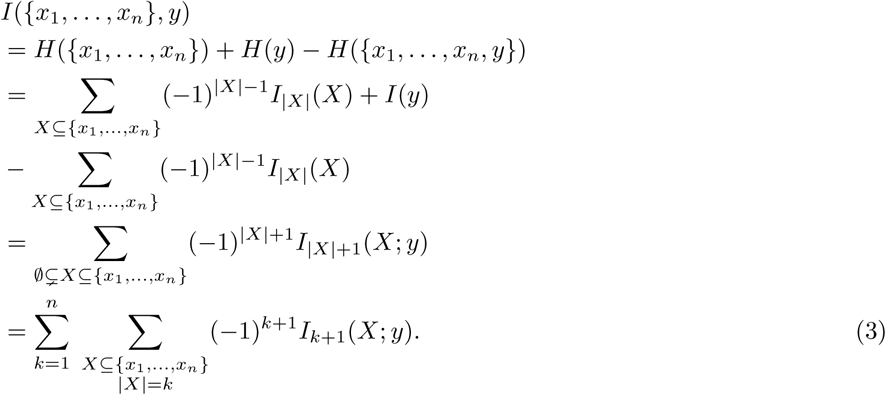

□

#### A.1 Degree K Supervised Feature Selection Algorithm

It is clear that if *X* ⊆*Y*, then *I*_*|X|*_(*X*) ≥ *I*_*|Y|*_(*Y*). In a practical setting it is also reasonable to believe that as *k* gets larger, the terms in the series in (3) converge to 0. As mentioned earlier, the computational cost of the terms in the series above grows exponentially with respect to *k*. This suggests that a fast “degree *K*” approximation to the mutual information *I*({*x*_1_, *…, x*_*n*_}; *y*):

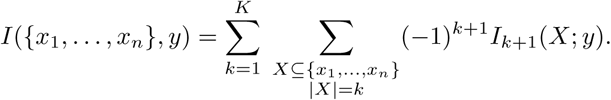

At each iteration of the greedy algorithm, our goal is to compute

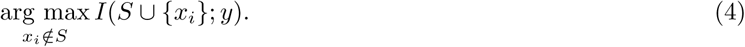

If we use the above degree *K* approximation, we only need to consider the terms involving *x*_*i*_ in the series expansion. Therefore, at each step, the degree *K* mRMR algorithm selects the next feature to add to *S* as:

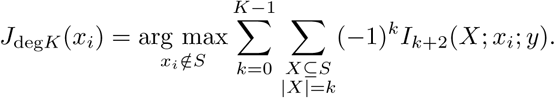

Notice that for *K* = 2, this simplifies to

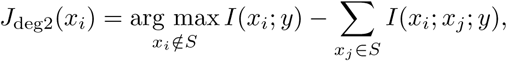

the CIFE objective in [12]. These functions *J*_deg*K*_ serve as the objective function *J* in the greedy search algorithm (Algorithm 1).

#### A.2 Degree K Unsupervised Feature Selection Algorithm

In the unsupervised setting, we are trying to solve

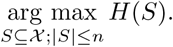

This involves approximating *n*-dimensional joint probability distributions, which necessitates the need for lower-degree approximations.

We use Corollary 2 to expand *H*(*S*) and write

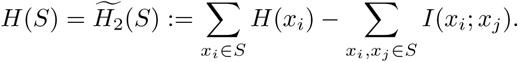

If we assume that there are no higher-degree interactions, we get a low degree approximation for *H*(*S*).

##### Corollary 4

*If I*_*|U|*_(*U*) = 0 *whenever |U |* ≥ 3, *then*

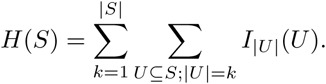

□

Using the greedy search (Algorithm 1), at each step, we identify

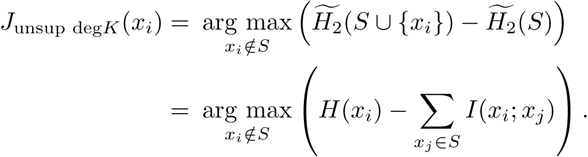

### B Guarantees for Greedy Search

The greedy search algorithm (Algorithm 1) is a necessary compromise to avoid the computation cost of an exhaustive search over all subsets of size at most *n*. However, there are some cases where such an approach is too shortsighted, even when we use the ideal objective function *J*_cmi_(*x*) = *I*(*S* ∪ {*x*}; *y*).

*Example* 1. Let *b*_1_, *b*_2_ be random variables chosen from {0, 1} independently with uniform probability. Let *b*_3_ be a random variable (independent of *b*_1_, *b*_2_) such that 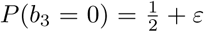 and 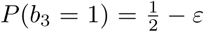 where 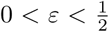. Finally, let *x*_1_ = *b*_1_, *x*_2_ = *b*_2_, *x*_3_ = *b*_1_ ⊕ *b*_2_ ⊕ *b*_3_, and *y* = *b*_1_ ⊕ *b*_2_. It is clear that *I*(*x*_1_; *y*) = *I*(*x*_2_; *y*) = 0 and *I*(*x*_3_; *y*) = 1 *– H*(*b*_3_). The optimal 2-element subset *S* of {*x*_1_, *x*_2_, *x*_3_} that maximizes *I*(*S*; *y*) is obviously {*x*_1_, *x*_2_}, but a greedy approach will always choose *x*_3_ at the first step.

In this section, we describe conditions under which we can give performance guarantees for the greedy search algorithm. These results rely heavily on results for the maximization of submodular functions. A function *f* defined on sets is submodular if it satisfies the diminishing returns property: *f* (*T* + {*x*}) *– f* (*T*) ≤ *f* (*S* + {*x*}) *– f* (*S*) whenever *S* ⊆*T*. The seminal paper on this topic by [27] proves the following result.

#### Theorem 5

([27]). *Consider the maximization problem*

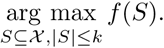

*If f is a non-decreasing submodular function, the greedy algorithm yields a (*1 *–* 1*/e)-approximation. In other words, if S*_*o*_ *is an optimal set and S*_*g*_ *is the set found via a greedy search, then*

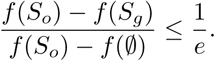

#### Submodularity of Entropy and its Degree 2 Approximation

##### Proposition 6

*The entropy function H*: 2^𝒳^→ℝ *is submodular.*

*Proof.* We want to show *H*(*T* ∪ {*x*}) *– H*(*T*) ≤ *H*(*S* ∪ {*x*}) *– H*(*S*) whenever *S* ⊆ *T*. Let *T’* = *T – S*. We have *I*(*T’*; *x|S*) ≥ 0 because (conditional) mutual information is always non-negative. This gives us

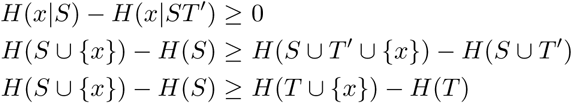

□

##### Proposition 7

*The degree 2 approximation* 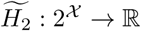 *of H, given by* 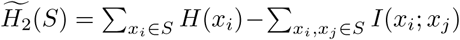 *is submodular.*

*Proof.* Note that 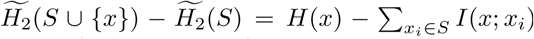 Recall that *I*(*·*; *·*) is non-negative. Clearly, 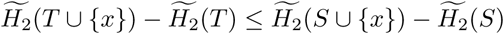 whenever *S* ⊆ *T*.

□

#### Submodularity of Mutual Information and its Degree 2 Approximation

In general, mutual information is not submodular. As we saw in Example 1, the “value” of a feature is not necessarily diminishing. In that particular example we had *I*(*x*_2_; *y*) *– I*(∅; *y*) = 0, but *I*({*x*_1_, *x*_2_}; *y*) *– I*(*x*_1_; *y*) = 1; crudely put, a greedy algorithm only appreciates the value of *x*_2_ when it already knows *x*_1_. This specific phenomenon is a direct consequence of synergy and we explain this relation rigorously in this section.

##### Proposition 8

*If for all x* ∉ *T* ⊇ *S, we have I*(*T –S*; *x*; *y |S*) ≥ 0, *then the set function f*: 2^𝒳^ →ℝ *given by f* (*S*) = *I*(*S*; *y*) *is submodular.*

*Proof.* As before, let *T*^*′*^ = *T – S*. We have *I*(*T*^*′*^; *x*; *y|S*) = *I*(*T*^*′*^; *x*; *y*) - *I* (*S*; *T* ^*′*^; *x*; *y*). We can now rewrite *I* (*T – S*; *x*; *y|S*) ≥ 0 as *I*(*S*; *T*^*′*^; *x*; *y*) *– I*(*T*^*′*^; *x*; *y*) ≤ 0. Adding many terms to each side of the inequality, we get

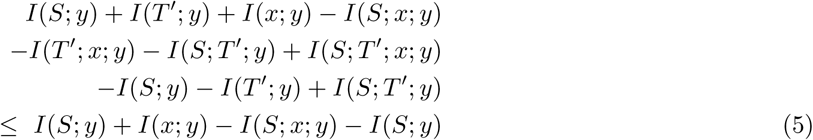

Therefore,

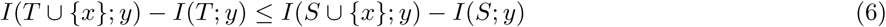

□

The analogous result for the degree approximation 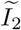 of *I* given by 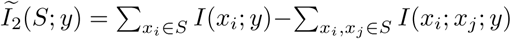 (which is the function that the CIFE algorithm maximizes) is a little easier to describe, since we ignore higher order interactions. We only need to assume that there is no two-way synergy.

##### Proposition 9

*If for all x*_*i*_, *x*_*j*_, *we have I*(*x*_*i*_; *x*_*j*_; *y*) ≥ 0, *then the set function f*: 2^𝒳^ → ℝ *given by* 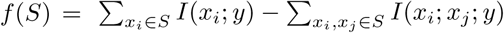 *is submodular.*

*Proof.* Note that 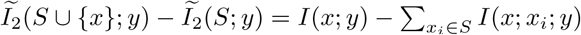. Because we have assumed *I*(*x*_*i*_; *x*_*j*_; *y*) ≥ 0 for all *x*_*i*_, *x*_*j*_, we have *f* (*T* ∪ {*x*}) *– f* (*T*) ≤ *f* (*S* ∪ {*x*}) *– f* (*S*) whenever *S* ⊆ *T*.

□

#### Guarantees for Feature Selection

It is well known that *I*(*·*; *y*) and *H*(*·*) are non-decreasing set functions. We immediately have the following corollaries.

##### Corollary 10

*The greedy search algorithm to maximize H*(*S*) *over S* ⊆ 𝒳 *with |S|* ≤ *k for some user-specified k yields a* (1 *–* 1*/e*)*-approximation (i.e., H*(*S*_*greedy*_) ≥ (1 *–* 1*/e*)*H*(*S*_*optimal*_).

*Proof.* Follows by Theorem 5 and Proposition 6.

##### Corollary 11

*If for all x* ∉ *T* ⊇ *S, we have I*(*T – S*; *x*; *y|S*) ≥ 0, *then the greedy search algorithm to maximize I*(*S*; *y*) *over S* ⊆ 𝒳 *with |S|* ≤ *k for some user-specified k yields a* (1 *–* 1*/e*)*-approximation (i.e., I*(*S*_*greedy*_; *y*) ≥ (1 *–* 1*/e*)*I*(*S*_*optimal*_; *y*).

*Proof.* Follows by Theorem 5 and Proposition 8.

Unfortunately, our approximations 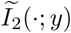 and 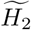 of *I*(*·*; *y*) and *H* functions are not non-decreasing, even though *I*(·; *y*) and *H* themselves are. This leaves us with a slightly weaker result for optimizing 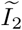 and 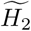. The following is a generalization of Theorem 5.

##### Theorem 12

([27]). *Consider the maximization problem*

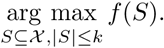

*If f is a submodular function and f* (*S* ∪ {*x*}) *– f* (*S*) ≥ *–θ for all S* ⊆ 𝒳 *and x* ∈ 𝒳, *we have*

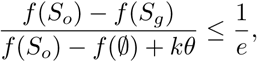

*where S*_*o*_ *is an optimal set and S*_*g*_ *is the set found via a greedy search.*

##### Corollary 13

*If* 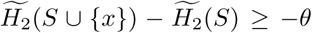 *for all S* ⊆ 𝒳 *and x* ∈ 𝒳, *then the greedy solution S*_*g*_ ⊆ 𝒳 *to* 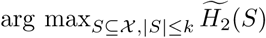 *relates to the optimal solution S*_*o*_ *as follows:*

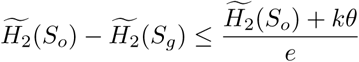

##### Corollary 14

*If I*(*x*_*i*_; *x*_*j*_; *y*) ≥ 0 *for all x*_*i*_, *x*_*j*_ ∈ 𝒳 *and* 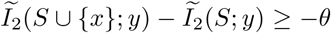 *for all S* ⊆ 𝒳 *and x* ∈ 𝒳, *then the greedy solution S*_*g*_ ⊆ 𝒳 *(i.e., the output of the CIFE algorithm) to* 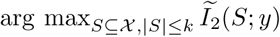 *relates to the optimal solution S*_*o*_ *as follows:*

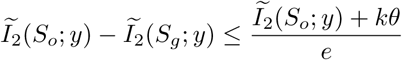

In practical terms, we can interpret Corollary 14 as follows: if we know that shared redundancy between any pair of features outweigh synergies and that our degree 2 approximation to mutual information is monotone, then the greedy algorithm will output an 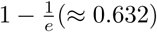 approximation (i.e., the score of the set returned by a greedy search will be at least 63.2% of the score of the optimal set). If we know that the degree 2 approximation is not monotone but can bound how much it might decrease by at any step, we can still recover a (weaker) guarantee.

### C Practical Implementation Issues

There are numerous issues that arise in the implementation of the information-theoretic algorithms in this paper. These issues warrant further discussion as they have a significant impact on the output of these algorithms. We present our approach to addressing these issues in this section. Optimizations that improve runtime but do not affect the output of the algorithm are presented under Efficient Algorithms.

#### C.1 Binning Technique

All our discussion thus far has assumed that the *x*_*i*_’s are discrete random variables. When working with continuous data, it is common to discretize the data prior to running information theoretic methods. Although the datasets we have used are inherently discrete (with entries in ℤ_≥0_), there is a need to limit the number of values that entries can take. To understand why this is so important, suppose that the *x*_*i*_’s take integer values from 0 to *k –* 1. In order to empirically compute *I*(*x*_*i*_; *x*_*j*_; *y*), we need to approximate *P* (*x*_*i*_, *x*_*j*_, *y*). Without any additional assumptions on the joint probability distribution, we are left to approximate this distribution by counting occurrences of each possible combination, of which there are *k*^3^. If our dataset has *n* observations, we need *n* ≫ *k*^3^ to have a reasonable approximation of *I*(*x*_*i*_; *x*_*j*_; *y*). It is also important not to collapse too many values together as that would result in very poor resolution stifling our ability to observe significant differences. We recommend targeting *k* = 5 discrete values as a reasonable default.

The process of choosing bins has a surprisingly large impact on the performance of the algorithms in this paper. For single cell RNA-sequencing, we propose a recursive quantile transformation that adjusts for the sparsity of the data. To motivate the need for our modification to the usual quantile transform, suppose we would like *k* = 5 bins for each feature. Because scRNA-seq datasets have a large number of zero entries (in some cases *>* 90% of entries are zero), 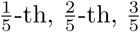, and 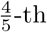 quantiles could all be 0. This leaves us with only two meaningful bins. To avoid this problem, we recommend a recursive strategy described in Algorithm 2 for selecting right endpoints for *k* bins. On our experiments, we perform this transformation on each feature individually to account for the different distributions of values across features.

##### Algorithm 2 Recursive-quantile transform

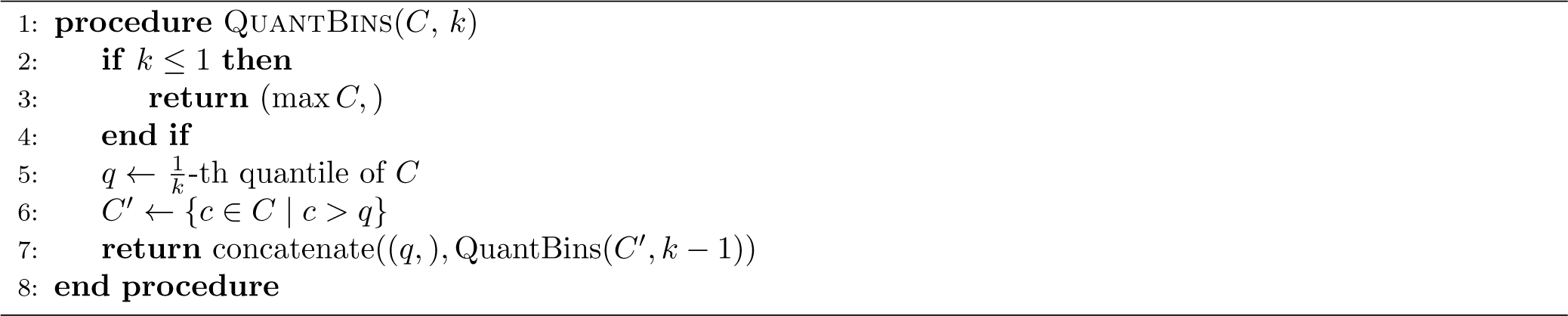

#### C.2 Multiclass vs Binary Feature Selection Methods

The mutual information based feature selection methods are approximations of arg max_|*S*|≤*n*_ *I*(*S; y*). Unlike many of the differential expression methods, the mutual information based algorithms allow for *y* to take more than two values (i.e., they are *multiclass* rather than *binary* methods). This, in theory, allows us to select features at once, rather than having to select them for every class label (one-vs-rest) or every pair of class labels (one-vs-one).

However, as we mentioned in the Results section, it is also possible to run these algorithms as binary methods by letting 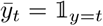 and treating 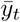 as the class labels. This gives us the opportunity both to compare binary differential expression methods to binary mutual information based methods and to evaluate strengths and weaknesses of multiclass methods vis-a-vis binary methods.

We saw that for almost all datasets (the only exception being the Zheng dataset for a small number of markers), turning the mutual information based methods into binary methods led to a significant improvement in performance. We describe some possible ways in which these differences in performance can be explained. Note that unlike in the Analysis of Existing Methods section, this is not an exhaustive list of shortcomings. Instead, these examples demonstrate ways in which these approaches differ.

Sometimes features can be informative about multiple target labels. Unless the feature selection algorithm used is inherently multi-class, we cannot detect features that may have a weak signal about any particular target label but a strong signal that helps discern between sets of target labels.

*Example* 2. Let *b*_1_, *b*_2_ be random variables chosen from {0, 1} independently with uniform probability. Let *y* = *b*_1_*b*_2_ and let 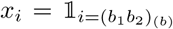 for *i* ∈ {0, 1, *…*, 3}. Here the subscript (*b*) denotes that we are looking at the binary representation of the number. For example,

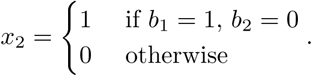

Finally, let *x*_4_ = *b*_1_ and *x*_5_ = *b*_2_.

It is easy to see now that *I*(*x*_4_; *y*) = 1 = *I*(*x*_5_; *y*). However, for *i* ≤ 3 we have 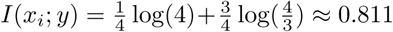. That is to say, multiclass methods will correctly identify *x*_4_ and *x*_5_ as being very informative about *y*.

On the other hand, if our class labels were binary, for *i* ≤ 3 we have 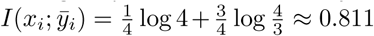.However,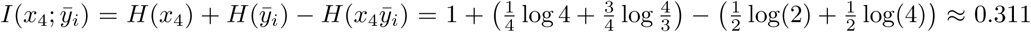.A binary method would therefore choose *x*_0_, *…, x*_3_ before *x*_4_ and *x*_5_.

It must be noted that if we run a binary method on the example above, we will eventually choose {*x*_0_, *x*_1_, *x*_2_, *x*_3_}, which, together, are as informative as {*x*_4_, *x*_5_}. The following modification demonstrates a situation in which this is not true.

*Example* 3. Let *c*_1_, *c*_2_, *…* be independent Bernoulli trials with probability *p* for some constant 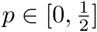 (these will introduce noise in *x*_0_, *x*_1_, *x*_2_, *x*_3_). In other words, ℙ(*c*_*i*_ = 1) = *p* and ℙ(*c*_*i*_ = 0) = 1 *– p*. Now follow the previous example, except let 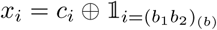 for *i* ∈ {0, 1, *…*, 3}. Recall ⊕ is the XOR operator: *a* ⊕ *b* = *a* if *b* = 0 and *a* ⊕ *b* = *¬a* if *b* = 1.

It is clear that for small values of *p*, a binary method will still choose {*x*_0_, *x*_1_, *x*_2_, *x*_3_} instead of {*x*_4_, *x*_5_}, even though *y* is a deterministic function of *x*_4_ and *x*_5_ but not of *x*_0_, *…, x*_3_. With a sufficiently large value of *p* (i.e., where there is enough noise in *x*_0_, *…, x*_3_), even a binary method will choose *x*_4_ and *x*_5_

Recall that objectives like CIFE and JMI only consider low-degree interactions between features. Therefore, it is easy to under/over-estimate redundancies. By measuring relevance (*I*(*x*_*i*_; *y*) term) and redundancies (*I*(*x*_*i*_; *x*_*j*_; *y*) terms) only in the context of a specific class label, binary methods are less vulnerable to accumulating errors in approximations.

We demonstrate this idea with two example. In the first example, we consider a univariate feature selection method (e.g., MIM) that does not penalize any redundancy (i.e., it underestimates redundancy) and outline a case where the binary version outperforms the multiclass version. In the second example, we demonstrate a situation in which a bivariate feature selection method (e.g., CIFE) overpenalizes redundancy in the multiclass version and the binary version yields a better selection of features.

*Example* 4. Let *b*_1_, *b*_2_, *b*_3_ be random variables chosen from {0, 1} independently with uniform probability. Let *y* = *b*_1_*b*_2_*b*_3_ and let 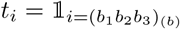 for *i* ∈ {0, 1, *…*, 7}. We will use the *t*_*i*_’s to define features. Let *x*_*i*_ = *t*_0_*t*_1_*t*_2_*t*_*i*_ for *i* ∈ {3, *…*, 7}. Let *x*_8_ = *t*_4_*t*_5_ and *x*_9_ = *t*_6_*t*_7_. A univariate multiclass method (e.g., MIM) will associate a higher score to *x*_3_, …, *x*_7_ because they are individually more informative about *y* than *x*_8_ and *x*_9_. However, a binary method (when considering the class label *y* = 100 or *y* = 101) will assign the same scores to *x*_8_, *x*_4_, and *x*_5_. Similarly, it would assign the same scores to *x*_9_, *x*_6_, and *x*_7_ when considering *y* = 110 and *y* = 111). To construct an example where the binary method will definitely pick *x*_8_ and *x*_9_ while the multiclass method won’t, we can add noise (as in Example 3) to features *x*_3_, …, *x*_7_.

In the previous example, we saw how binary methods, in searching for information specific to individual class labels, are able to avoid traps of redundant information. We now demonstrate an example in which binary methods avoid overpenalizing redundancy.

*Example* 5. Let *b*_1_, *b*_2_ be random variables chosen from {0, 1} independently with uniform probability. Let *c*_2_, *c*_3_ be independent Bernoulli trials with probability 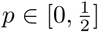. Let *y* = *b*_1_*b*_2_ and let 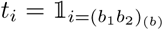 for *i* ∈ {0, 1, *…*, 3}. We will use the *t*_*i*_’s to define features. Let *x*_1_ = *t*_0_*t*_1_*t*_2_, *x*_2_ = *t*_0_*t*_1_*t*_3_, *x*_3_ = *t*_2_ ⊕ *c*_2_, and *x*_4_ = *t*_3_ ⊕ *c*_3_.

If we use a bivariate multiclass method (e.g., CIFE), without loss of generality we select *x*_1_ at the first iteration (analogous argument if we selected *x*_2_ first). Because *I*(*x*_2_; *x*_1_; *y*) is significant, the penalty on *x*_2_ (with CIFE, say) would be large enough that we would pick *x*_4_ instead of *x*_2_, which is more informative. The binary version of CIFE would not have this problem.

We posit that the Paul dataset, with its high cluster resolution (numerous clusters that cannot be separated easily) resembles Example 5 and believe that this (at least partially) explains the stark contrast in the reclassification accuracy between the multiclass and binary algorithms (especially CIFE).

### D Efficient Algorithms

#### D.1 Computing Entropy for Sparse Matrices

In this section, we outline a fast computation of entropy for sparse matrices. As a result of Proposition 1, this algorithm can be used to efficiently compute various information-theoretic objective functions for feature selection (especially low-dimension approximations, such as CIFE and JMI). In this section, we will use zero-indexing rather than one-indexing (i.e., the first element of an array *A* is *A*[0]).

Compressed Sparse Column (CSC) is a sparse matrix format that is optimized for column operations (e.g., slicing by columns, iterating over non-zero entries in a given column, etc.). It is defined as follows. For an *n ×m* matrix *M*, the CSC format stores three arrays:

- *X*, the values of the non-zero entries stored in column major order.
- *R*, the row index of the non-zero entries in *X* (i.e., *X* and *R* have the same length).
- *C*, an array of length *m* + 1 where *C*[*j* + 1] *– C*[*j*] is the number of non-zero entries in column *j* and *C*[0] = 0.

In other words, for any *s* such that *C*[*j*] ≤ *s < C*[*j* + 1], we have *X*[*s*] is the value of a non-zero entry in row *R*[*s*] and column *j*.

Assume all values in *X* come from {0, 1, *…, k –* 1} (i.e., there are *k* distinct values in *X*). To efficiently compute the (empirical) entropy of an individual column *j* of an *n × m* matrix *M*, we follow Algorithm 3.

##### Algorithm 3 Entropy of a Single Column

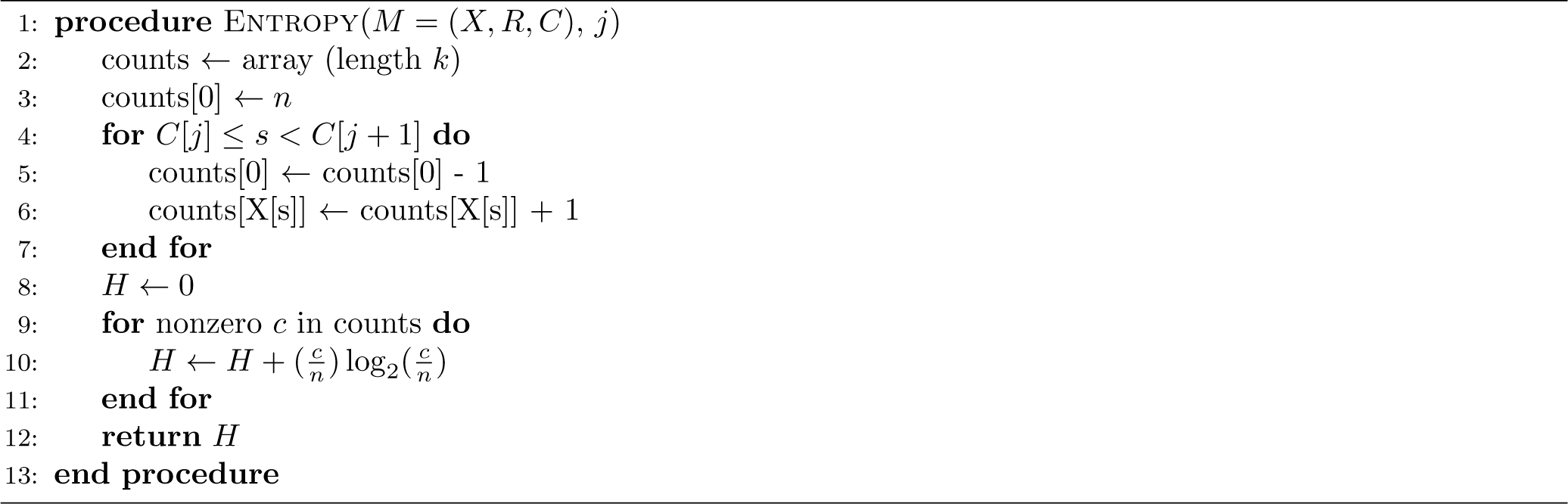

We often need to compute the entropy of two or more columns. A trivial way to overcome this problem is to combine multiple columns into one column. One way to combine *r* columns is to write each entry as an *r*-tuple. This combined column will retain the sparse structure as the non-zero entries will correspond to rows in which at least one entry was non-zero.^3^

However, the process of computing the tuple representation of multiple can sometimes be repetitive. For example, to compute arg 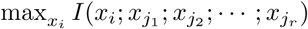, we need to compute 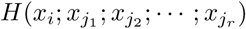 for all *x*_*i*_ ∈ 𝒳. In such cases, it make sense to precompute the tuple representation of 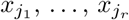.

However, in most cases, the column of class labels *y* tends not to be a sparse column. Combining *y* into the tuple representation means that most entries in the column become non-zero. This voids the benefits of exploiting the sparse structure of *M*. We can address these concerns by precomputing some information and vectorizing the entropy computation as in Algorithm 4. The ENTROPYWRT (“entropy with respect to”) takes a list of columns 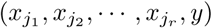 and returns the vector 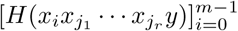. Pseudocode for ENTROPYWRT when the *y* column is not in the list of columns can be written with a trivial modification to Algorithm 4. Here we assume *y* has entries in 0, 1, *…, n*_labels_ – 1.

The time complexity of ENTROPYWRT is *O*(*n* + *mp* + *q* + *mk*^*r*+1^) where *p* is the number of rows with non-zero entries in columns *j*_1_, *j*_2_, …, *j*_*r*_, and *q* is the number of non-zero entries in *M*. Typically, ENTROPYWRT will be called numerous times, in which case ycounts should be precomputed once, rather than on each call to ENTROPYWRT. In this case, the time complexity is *O*(*mp*+*q* +*mk*^*r*+1^). Also note that when we use ENTROPYWRT, we will typically fix *k* beforehand (depending of discretization strategy) to be about 5. For CIFE and JMI algorithms, we have *r* = 2 because we will only ever compute the joint entropy of three columns or less. In these cases, the time complexity is *O*(*mp* + *q*).

#### D.2 Efficient Optimization of CIFE and JMI

The CIFE and JMI algorithms compute arg 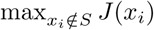 at each iteration. If done naively, computing *J* (*x*_*i*_) can take longer as *S* grows because 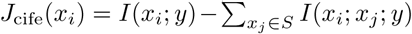 and 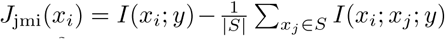 have *|S|* + 1 terms. The computational complexity to select *t* features is *O*(*mt*^2^*f*) where *f* is the time to compute *I*(*x*_*i*_; *x*_*j*_; *y*) and is a function of *n, k*, and sparsity parameters of columns in *X*. A minor modification to vectorize this process can lower the complexity to *O*(*mtf*), as shown in Algorithm 5.

Note that *I*(*x*_*s*_; *x*_*j*_; *y*) = *I*(*x*_*j*_; *y*) + *H*(*x*_*s*_) *– H*(*x*_*s*_, *x*_*j*_) *– H*(*x*_*s*_, *y*) + *H*(*x*_*s*_, *x*_*j*_, *y*). Since *I*(*x*_*j*_; *y*) was computed as basescore, and *H*(*x*_*s*_) *– H*(*x*_*s*_; *y*) is constant with respect to *j*, the only two calls to ENTROPYWRT are for *H*(*x*_*s*_, *x*_*j*_) and *H*(*x*_*s*_, *x*_*j*_, *y*).

Under certain conditions, this process can be made even faster with a strategy adapted from the fast algorithm for CMIM in [28]. In the aforementioned paper, the authors propose a objective function that can be rewritten as 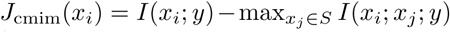. They improve the efficiency of the naive algorithm by updating *J*_cmim_(*x*_*i*_) (i.e., checking if the penalty term has increased since more features were selected into *S*) only *J* (*x*_*i*_) *> J* (*x*_*j*_) for all *j < i*. This prevents some unnecessary evaluations of *I*(*x*_*i*_; *x*_*j*_; *y*). This trick hinges on the monotonicity of 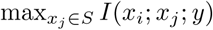 with respect to *S*. In our case Σ*x*_*j*_ ∈ *SI*(*x*_*i*_; *x*_*j*_; *y*) is monotone if *I*(*x*_*i*_; *x*_*j*_; *y*) ≥ 0. If we do make this assumption (as we did in Proposition 9), we could make a similar optimization. In fact, we believe such an optimization would be best implemented using a priority queue.

##### Algorithm 4 Entropy of Multiple Columns (all but one fixed)

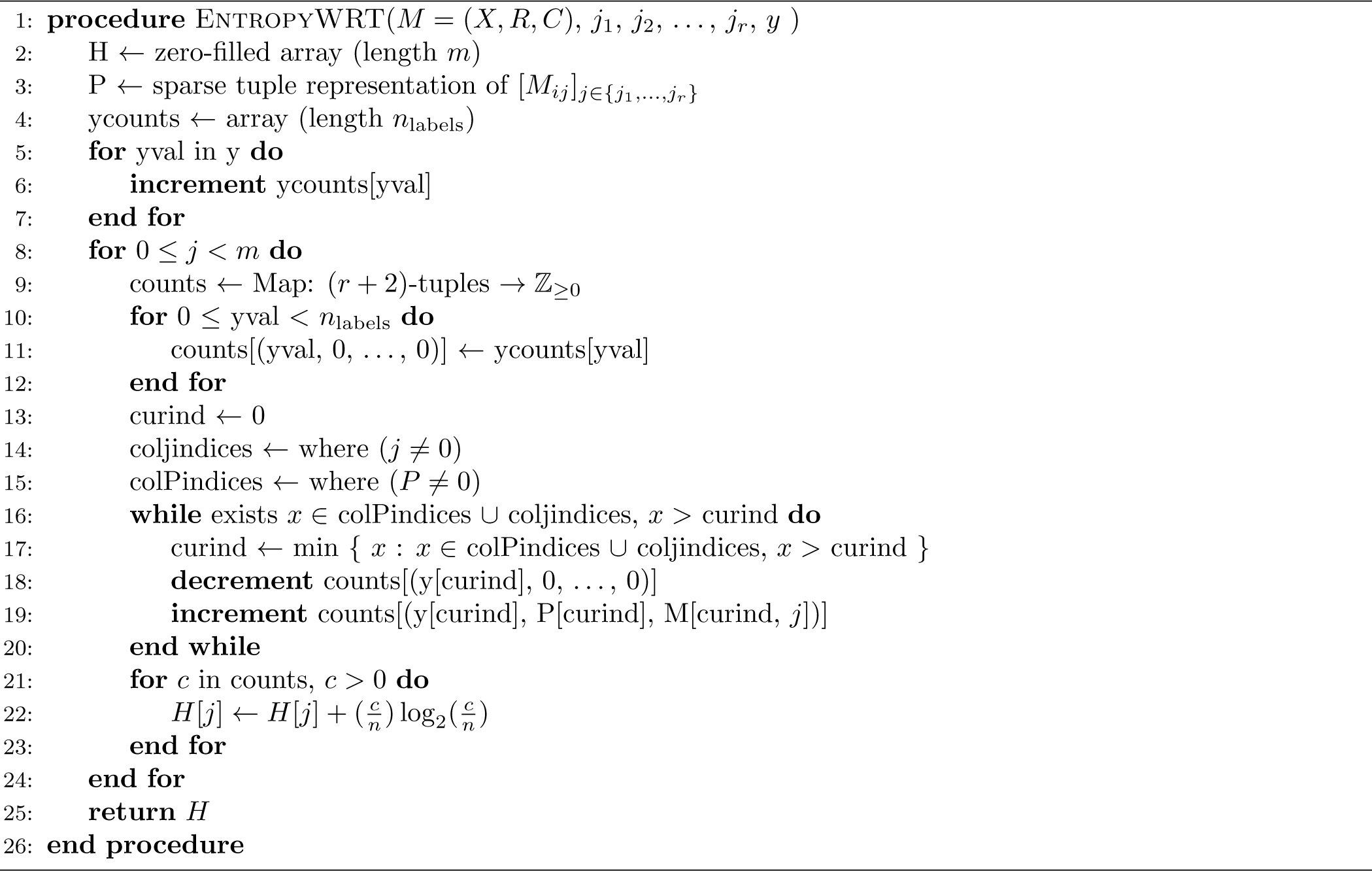

##### Algorithm 5 Vectorization of CIFE/JMI

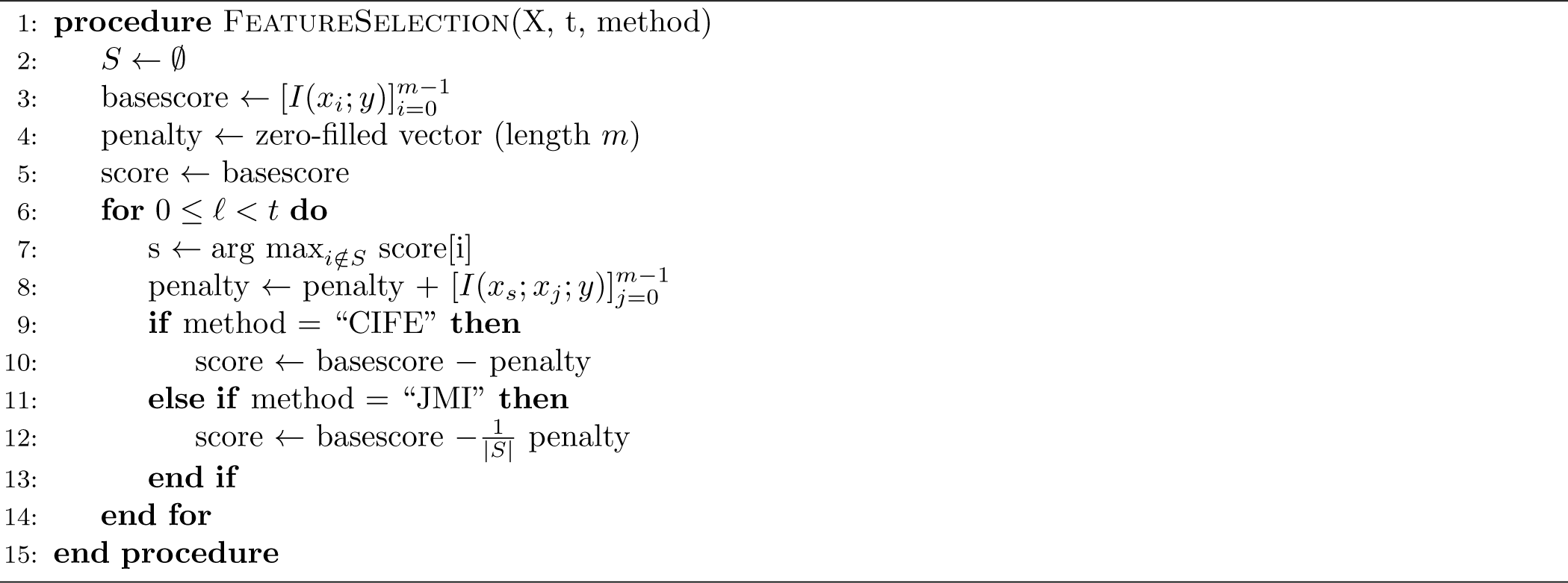

### E Analysis of Existing Methods

Most of these algorithms are affected by three broad theoretical weaknesses. We analyse these problems in the context of existing algorithms and present simple examples where they arise. Obviously, many assumptions from [20] that we discussed under Previous work are violated in these simple examples. The aim of these examples, however, is to present the theoretical weaknesses of existing algorithms in information-theoretic terms. This will motivate our approach to mutual information based feature selection^4^.

#### E.1 Diminishing penalty on redundancy

The mRMR algorithm in [21] is a prominent example of an objective that progressively diminishes the penalty on redundancy. Specifically,

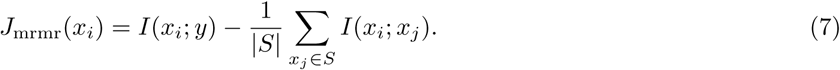

The JMI algorithm in [22] also has a factor of 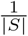 on its redundancy term:

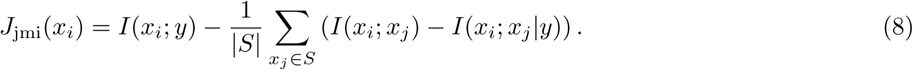

When *|S|* is large, the redundancy penalty is so small that these algorithms, when asked for a large set of features (but still considerably smaller than the total number of features), will yield similar outputs as MIM. We demonstrate this empirically in Tables 1, 2, 4, and 3.

In practice, the 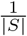 factor in algorithms such as mRMR and JMI serves two purposes. First, the 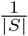 factor accounts for shared redundancy between multiple variables (see examples under Ignoring Higher-degree Interactions Between Features). Second, it acts to mitigate the accumulating noise in empirical estimations of *I*(*x*_*i*_; *x*_*j*_), because even if *x*_*i*_ and *x*_*j*_ are independent, the empirical estimation of their mutual information might be positive. In either case, the 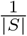 factor is a somewhat arbitrary approach that suppresses both signal and noise and, in doing so, impedes our ability to detect true redundancies.

*Example* 6. Let *b*_1_, *b*_2_, *…* be random variables chosen from {0, 1} independently with uniform probability. Let *x*_1_ = *b*_1_*b*_2_ … *b*_5_ (a binary concatenation of bits), *x*_2_ = *b*_6_*b*_7_ … *b*_10_, *x*_3_ = *b*_11_, and *x*_4_ = *b*_1_*b*_2_*b*_3_. Suppose *y* = *b*_1_*b*_2_ … *b*_11_. Algorithms such as mRMR and JMI would select *x*_1_ and *x*_2_ first. At this point, *J* (*x*_3_) = 1 and 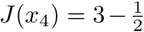·3 = 1.5. But clearly, *x*_4_ contributes no new information, while *x*_3_ does.

#### E.2 Penalizing Irrelevant Redundancy

Algorithms like mRMR and MIFE penalize all redundancy, even if the redundancy does provide any information about *y*. For example, if *x*_*i*_ and *x*_*j*_ are highly correlated, then it is possible that *I*(*x*_*i*_; *x*_*j*_) *>* max (*I*(*x*_*i*_; *y*), *I*(*x*_*j*_; *y*)). This is demonstrated in the following example.

*Example* 7. Let *b*_1_, *b*_2_, *…* be random variables chosen from {0, 1} independently with uniform probability. Let *x*_1_ = *b*_1_*b*_2_*b*_3_, *x*_2_ = *b*_1_*b*_2_*b*_4_, *x*_3_ = *b*_5_, and *y* = *b*_3_*b*_4_. Without loss of generality, assume the algorithm chooses *x*_1_ first. Now *I*(*x*_2_; *y*) = 1, but *I*(*x*_1_; *x*_2_) = 2. Such algorithms might choose *x*_3_ instead of *x*_2_, because *I*(*x*_1_; *x*_2_) is large, but the redundancy between *x*_1_ and *x*_2_ is not relevant to *y* and should not be penalized.

#### E.3 Ignoring Higher-degree Interactions Between Features

Features can say both more and less about the target labels together than the sum of their parts. All features covered by Brown, et al.’s framework (objectives of the form (2)) are unable to detect higher-order synergies and redundancies because they only detect two-way interactions between features.

*Example* 8 (Synergy). Let *b*_1_, *b*_2_, *…* be random variables chosen from {0, 1} independently with uniform probability. Let *x*_1_ = *b*_1_ and *x*_2_ = *b*_2_. If *y* = *b*_1_ ⊕ *b*_2_ (where ⊕ denotes the XOR operation on bits), then *I*(*x*_1_; *y*) = *I*(*x*_2_; *y*) = 0, but *I*(*x*_1_*x*_2_; *y*) = 1. This can be understood in terms of the multivariate mutual information *I*(*x*_1_; *x*_2_; *y*) = −1 (which we saw in the Series Expansion of Mutual Information section)^5^

*Example* 9 (Shared Redundancy). Let *b*_1_, *b*_2_, *…* be random variables chosen from {0, 1} independently with uniform probability. Let *x*_1_ = *b*_1_*b*_4_, *x*_2_ = *b*_2_*b*_4_, *x*_3_ = *b*_3_*b*_4_, *x*_4_ = *b*_5_, and *y* = *b*_1_*b*_2_*b*_3_*b*_4_. Without loss of generality, after two iterations, an algorithm of the form (2) will likely have *S* = {*x*_1_, *x*_2_}. Now, *I*({*x*_1_, *x*_2_, *x*_3_}, *y*) = 4 and *I*({*x*_1_, *x*_2_, *x*_4_}, *y*) = 3, but objectives of the form (2) penalize *x*_3_ twice for its redundant bit (*b*_4_).

### F Problems with mRMR

A false theorem in [21, *§*2.3] states that maximizing the mRMR objective in a greedy search is “equivalent” to maximizing the CMI objective in a greedy search. The authors of this paper believe that the reasonable interpretation of “equivalent” is that given the same input, the two algorithms will return the same output. In this case, their claim (and the proof thereof) is incorrect. We have already seen in Examples 6 and 7 cases where the mRMR objective makes different choices than the CMI objective. To understand why these examples are counterexamples to the claim in [21], we walk through their proof.

For notational convenience, we let 𝒳 be the set of all features, *S* = *S*_*m–*1_ be the set of *m –* 1 features selected after *m –* 1 steps, *D*(*x*_*i*_) = *I*(*x*_*i*_; *y*) be the *relevance* of feature *x*_*i*_ and 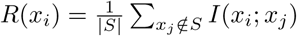 be the *redundancy* of feature *x*_*i*_. They definite a quantity *J*: *P*(𝒳) *→* ℝ and show that *I*(*S*_*m–*1_ ∪ {*x*_*i*_}; *y*) = *J* (*S*_*m–*1_, *x*_*i*_, *y*) *– J* (*S*_*m–*1_, *x*_*i*_). Fixing *S*_*m–*1_, they then show that the first term, *J* (*S*_*m–*1_, *x*_*i*_, *y*), is maximized when *D*(*x*_*i*_) = *I*(*x*_*i*_; *y*) is maximized, and the second term, *J* (*S*_*m-*1_, *x*_*i*_) is minimized when 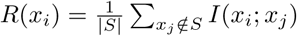 is minimized. They then falsely conclude that

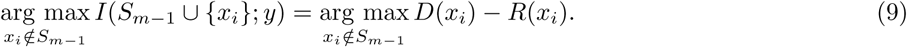

### Footnotes

1 We used the latest developmental version (commit 80e635d2b78) available at the time as this revision fixed a bug that prevented us from using scanpy during *K*-fold cross validation.

2 Brown et al. incorrectly claim IGFS is equilvalent to CIFE, it is, in fact, equivalent to JMI.

3 In practice, is can be faster to represent (*a*_0_, *a*_1_, …, *a*_*r−*1_) as *a*_0_ + *a*_1_*k* + + *a*_*r–*1_*k*^*r–*1^ where *k* = max_*i*_(*a*_*i*_) + 1, but we will avoid this conversion for the sake of readability. See our source code for such an implementation.

4 It should be noted, however, that no algorithm can truly overcome all the weaknesses below for that will require being able to infer a joint probability distribution over a large number of random variables which is meaningless without an enormous number of samples and also computationally intractable.

5 In cases of extreme synergy, such as the current example (where individual feature provide no information on their own) even the *J*_cmi_ objective will fail to choose an optimal set of features as a result of the shortsightedness of a greedy approach. In less extreme cases, it is still beneficial for the objective function to recognize such synergy).

## Notes

https://github.com/umangv/picturedrocks

https://github.com/umangv/picturedrocksbenchmarks

